# Mitochondrial-derived compartments buffer outer membrane protein load during acute mitochondrial adaptation

**DOI:** 10.64898/2026.06.09.731211

**Authors:** Bo J. Price, Sai Sangeetha Balasubramaniam, Adam L. Hughes

**Affiliations:** Department of Biochemistry, University of Utah School of Medicine, Salt Lake City, UT 84112, USA

## Abstract

Cells undergoing metabolic transitions rapidly remodel mitochondria through coordinated expansion and reorganization of the mitochondrial proteome. How the outer mitochondrial membrane (OMM) accommodates acute increases in newly synthesized proteins before organelle adaptation is complete remains poorly understood. Here we show that mitochondrial-derived compartments (MDCs), multilamellar domains that form from the OMM and selectively sequester OMM-associated cargo, arise during metabolic perturbations associated with acute mitochondrial biogenesis, including glucose restriction, carbon-source switching, and salt stress. In these situations, MDC formation requires the energy-sensing kinase Snf1 and derepression of the transcriptional repressor Mig1, linking MDC induction to transcriptional programs that increase mitochondrial protein expression. Activation of mitochondrial biogenesis in the absence of metabolic changes is sufficient to trigger MDCs, whereas disruption of mitochondrial protein targeting and import prevents MDC formation and causes mislocalization of outer membrane cargos. Together, these findings, combined with previous observations that MDCs are induced by hydrophobic protein overexpression, mistargeting, and metabolic perturbations, support an emerging model in which MDCs function as adaptive outer-membrane remodeling domains that buffer outer membrane protein load during mitochondrial adaptation.

## Introduction

Cells frequently undergo metabolic transitions that require mitochondria to rapidly remodel their composition and function to meet new energetic demands (Nunnari & Suomalainen, 2012; Friedman & Nunnari, 2014; Wai, 2024). During these adaptations, mitochondria adjust their proteome, membrane composition, and metabolic capacity to support new functional states. Multiple mechanisms have been discovered that coordinate these changes. For example, mitochondrial dynamics reshape the network to match metabolic conditions (Westermann, 2012; Youle & Van Der Bliek, 2012; Wai, 2024), mitophagy and vesicular trafficking pathways remove damaged or superfluous components (Palikaras *et al*, 2015; Pickles *et al*, 2018; Poillet-Perez & White, 2021), and transcriptional programs stimulate mitochondrial biogenesis to expand respiratory capacity (Kayikci & Nielsen, 2015; Piantadosi & Suliman, 2012; Martinez-Ortiz *et al*, 2019; Wang *et al*, 2024). In parallel, mitochondrial protein import and quality-control systems—including membrane potential–dependent import pathways and surveillance mechanisms that monitor mislocalized or unimported precursors—help maintain mitochondrial proteostasis during these transitions (Gebert *et al*, 2011; Kim *et al*, 2024; Wiedemann & Pfanner, 2017; Weidberg & Amon, 2018). Together, these processes remodel mitochondrial structure and composition as cells adapt to changing metabolic environments.

Despite these advances, acute metabolic transitions that are associated with increased mitochondrial biogenesis likely pose a fundamental challenge for mitochondria (Boos *et al*, 2019; Kim *et al*, 2024). These transitions often occur while mitochondrial function is compromised due to changes in metabolite availability. As mitochondrial biogenesis is rapidly induced, newly synthesized mitochondrial proteins are delivered to an organelle that is still adapting (Pfanner *et al*, 2019; Pastor *et al*, 2009; Lackner, 2014). This influx may transiently exceed the capacity of membrane organization, targeting, or assembly pathways needed to accommodate it. Such transitional states may place unique demands on the outer mitochondrial membrane (OMM), the first landing site for many newly synthesized mitochondrial proteins during biogenesis. How mitochondria manage this transient protein surge before the organelle is fully adapted remains poorly understood.

The mitochondrial-derived compartment (MDC) pathway is a recently described remodeling response of the mitochondrial outer membrane that may be uniquely positioned to buffer acute increases in OMM protein load. MDCs are multilamellar membrane domains derived from the OMM that selectively capture and sequester hydrophobic outer-membrane proteins away from the organelle (Hughes *et al*, 2016; Wilson *et al*, 2024b). Previous work has shown that MDC formation can be triggered by diverse metabolic perturbations that alter mitochondrial homeostasis, including changes in cellular amino acids, mitochondrial metabolites, and lipids (Schuler *et al*, 2021; Raghuram & Hughes, 2024; Xiao *et al*, 2024; Wong *et al*, 2025). MDCs can also form when hydrophobic proteins accumulate at mitochondria due to overexpression or mistargeting following failed insertion at another organelle (Wilson *et al*, 2024a). Recent work has further implicated Yme1-dependent proteolytic remodeling of mitochondrial lipid-transfer and membrane-architecture factors in enabling MDC biogenesis, supporting the idea that MDC formation requires coordinated remodeling of mitochondrial membrane state as well as cargo load (Balasubramaniam *et al*, 2026). Together, these observations suggest that MDC formation is associated both with mitochondrial metabolic perturbation and with increased hydrophobic membrane protein burden, although whether MDCs normally function during physiological mitochondrial adaptations remains unknown.

Here, while working to identify physiological metabolic alterations that induce MDC formation and thereby better define the role of these structures in cellular physiology, we found that conditions that stimulate acute mitochondrial biogenesis—including glucose restriction, carbon-source switching, and salt stress—robustly induce MDC formation in *Saccharomyces cerevisiae*. MDC induction in these contexts requires Snf1, which promotes MDC formation by relieving Mig1-mediated repression of mitochondrial biogenesis programs. Consistent with these data, we show that increased mitochondrial protein expression also occurs in previously identified metabolic MDC-inducing conditions, albeit in a Snf1-independent manner. Moreover, inducible activation of mitochondrial biogenesis through the transcriptional regulator Hap4, even in the absence of acute metabolic alterations, is sufficient to trigger MDC formation, whereas blocking delivery of hydrophobic proteins to mitochondria prevents MDC formation. Together with previous observations, these findings support a unifying model in which MDCs function as adaptive outer-membrane remodeling domains that are engaged when the OMM experiences elevated or perturbed protein load—a situation that arises in a variety of contexts, including protein mistargeting (Wilson *et al*, 2024a), changes in membrane lipids (Wong *et al*, 2025; Xiao *et al*, 2024), or during acute metabolic transitions that increase mitochondrial protein load while mitochondria are in a vulnerable metabolic state (current study; Hughes *et al*, 2016; Schuler *et al*, 2021).

## Results

### Physiological stresses that stimulate acute mitochondrial biogenesis induce MDC formation

Previous work from our laboratory showed that MDC formation can be triggered by metabolic stresses that perturb mitochondrial homeostasis, including changes in amino-acid availability or mitochondrial lipid balance, as well as by conditions that increase mitochondrial membrane protein load, such as forced overexpression or mistargeting of hydrophobic outer-membrane proteins (Hughes *et al*, 2016; Wilson *et al*, 2024a; Schuler *et al*, 2021; Raghuram & Hughes, 2024; Xiao *et al*, 2024; Wong *et al*, 2025). These observations suggested that MDCs arise when mitochondrial metabolic perturbation coincides with increased membrane protein burden. We therefore reasoned that acute metabolic transitions that alter mitochondrial metabolic state while also activating mitochondrial biogenesis might naturally generate these conditions. To test this idea, we examined metabolic stresses and transitions known to stimulate mitochondrial biogenesis and alter mitochondrial metabolism, including glucose restriction, abrupt carbon-source switching from fermentable to non-fermentable sugars, glycolytic inhibition, and salt stress (Coccetti *et al, 2018*; Martinez-Ortiz *et al*, 2019; Pastor *et al*, 2009; Posas *et al*, 2000; Kayikci & Nielsen, 2015).

We first tested whether glucose restriction, a physiological inducer of mitochondrial biogenesis, triggers MDC formation. MDCs were monitored by fluorescence microscopy as previously described (Schuler *et al*, 2021) using the outer-membrane marker Tom70-GFP together with the inner-membrane marker Tim50-mCherry, where MDCs appear as Tom70-positive domains that exclude Tim50 signal. Consistent with activation of mitochondrial biogenesis, several mitochondrial proteins—including the outer-membrane markers Om45 and Por1 as well as the matrix enzyme Cit1—displayed increased expression within one to two hours after glucose restriction (Fig. 1A, Fig. S1A). Strikingly, shifting cells from rich medium containing 2% glucose to rich medium without added glucose induced robust MDC formation, with 70–80% of cells forming Tom70-enriched MDCs within 2 hours (Fig. 1B-C).

**Figure 1:**
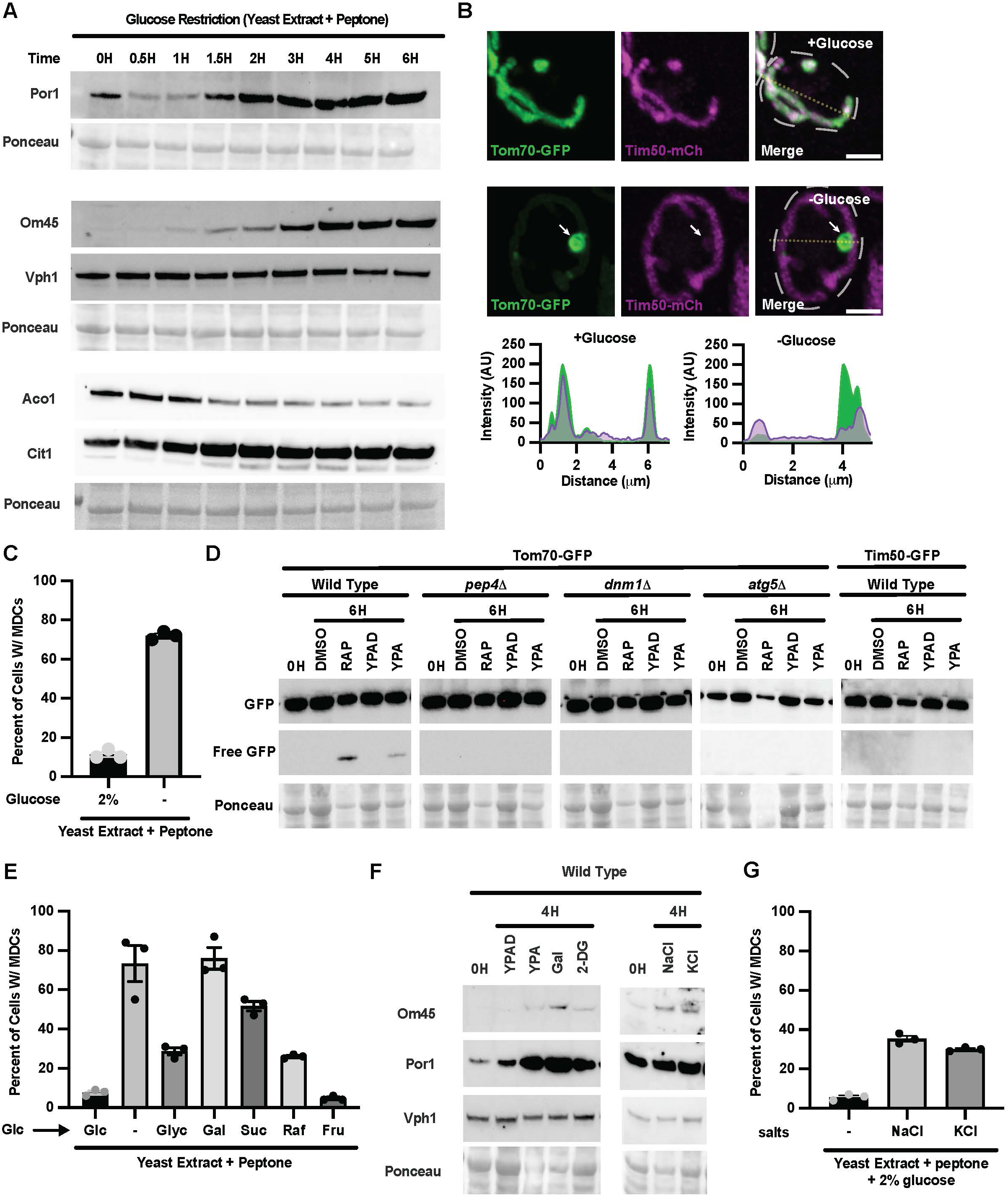
Metabolic stresses that stimulate acute mitochondrial biogenesis induce MDC formation. A. Time-course immunoblot of mitochondrial protein levels in wild-type yeast during glucose restriction (0-6 h). Por1, Om45, Aco1, and Cit1 were detected with the indicated antibodies; Vph1 and Ponceau staining serve as loading controls. B. Super-resolution images of wild-type cells grown in glucose-replete (YPAD) or glucose-restricted (YPA) media for 3 h. Cells express Tom70-GFP and Tim50-mCherry. Line-scan intensity profiles are shown below and correspond to the yellow dashed lines. The white arrow indicates an MDC. Scale bar, 2 μm. C. Quantification of the percentage of cells containing MDCs following 3 h glucose restriction. D. Immunoblot analysis of GFP processing after 6 h of the indicated treatments in wild-type*, pep4Δ*, *dnm1Δ*, and *atg5Δ* strains expressing Tom70-GFP, or wild-type cells expressing Tim50-GFP. Free GFP indicates proteolytic cleavage. Ponceau staining serves as a loading control. E. Quantification of MDC formation following a 3 h carbon-source shift. Cells were grown overnight in glucose (YPAD) and shifted to media containing the indicated carbon sources: glucose (Glc), no added carbon source (-), glycerol (Glyc), galactose (Gal), sucrose (Suc), raffinose (Raf), or fructose (Fru). F. Immunoblot analysis of mitochondrial protein levels following glucose restriction, carbon-source switching, glycolytic inhibition, or salt stress (0.8 M NaCl or 1 M KCl). Om45 and Por1 are shown; Vph1 and Ponceau staining serve as loading controls. G. Quantification of MDC formation after 3 h of salt stress (0.8 M NaCl or 1 M KCl). Quantification and statistics. For all MDC measurements, n = 3 independent experiments with 100 cells scored per replicate; bars show mean ± SEM.

Glucose restriction–induced MDCs appeared rapidly and excluded the inner-membrane marker Tim50-mCherry. Time-lapse imaging revealed that MDC biogenesis began with the formation of Tom70-GFP foci at mitochondria, which subsequently elongated and matured into multilamellar spherical structures within 0.5-1 h of treatment (Movie S1). Mature MDCs generated during glucose restriction reached approximately 0.70 microns in diameter on average, similar to previously reported MDCs induced by the mTOR inhibitor rapamycin (Fig. S1B). The cargo profile of glucose-restriction induced MDCs also resembled that of previously characterized MDCs triggered by treatment with rapamycin (Fig. S1C-D). In addition, glucose restriction–induced MDC cargos were delivered to the vacuole and degraded through the same autophagy– and vacuolar protease-dependent pathway used by MDCs formed under other stresses. GFP-processing assays showed that Tom70 and other established MDC cargos underwent degradation in a manner dependent on Dnm1-mediated fission, Atg5-dependent autophagy, and the vacuolar protease Pep4, whereas the inner-membrane protein Tim50 was not degraded (Fig. 1D, Fig. S1E) (Hughes *et al*, 2016; Schuler *et al*, 2021). Together, these observations indicate that glucose restriction is a physiological trigger for MDC-dependent OMM remodeling and removal of outer-membrane content.

We next asked whether other perturbations of glycolytic metabolism similarly induce MDC formation. Shifting cells from glucose to alternative carbon sources produced variable levels of MDC induction (Fig. 1E). These differences largely correlated with the degree of metabolic reprogramming required: switching to carbon sources that remain primarily fermentative, such as fructose, failed to induce MDCs, whereas carbon sources that are non-fermentable or require transcriptional activation of alternative metabolic programs, such as galactose through the GAL network, produced strong MDC responses (Nehlin *et al*, 1991; Platt & Reece, 1998). Inhibition of glycolysis with the starvation mimic 2-deoxy-D-glucose (2-DG) also induced MDC formation at intermediate concentrations, while higher concentrations failed to induce MDCs (Fig. S1F). These findings suggest that MDC biogenesis is activated when glycolytic metabolism is altered.

Because glucose restriction and carbon switching from fermentable to non-fermentable sugars stimulate mitochondrial biogenesis, we next tested whether another stress that triggers mitochondrial metabolic remodeling could also induce MDC formation. Salt and osmotic stress impair central carbon metabolism and are well known to activate Sucrose Non-Fermenting 1 (Snf1) kinase, a central regulator of mitochondrial biogenesis (Pastor *et al*, 2009; Posas *et al*, 2000). Consistent with this known stimulatory effect on mitochondrial biogenesis, exposure to 0.8 M NaCl increased mitochondrial protein expression with kinetics similar to glucose restriction, carbon switching, and 2-DG treatments (Fig. 1F). Both NaCl and KCl induced MDC formation, with approximately 35% of cells having MDCs after 3 hours (Fig. 1G). Taken together, these results indicate that MDCs are induced by metabolic stresses that stimulate acute mitochondrial biogenesis while altering mitochondrial metabolism. Glucose restriction, carbon-source transitions, glycolytic inhibition, and salt stress each perturb metabolism and increase mitochondrial protein expression, suggesting that MDC formation accompanies specific metabolic transitions that drive mitochondrial remodeling.

### MDC formation during glucose restriction requires trace glucose but is independent of amino-acid availability

To better understand how glucose restriction induces MDC formation, we next investigated the metabolic requirements underlying this response. Our previous work showed that several MDC-inducing stresses—including mTOR inhibition, impaired vacuolar function, and translation perturbation—stimulated MDC formation by elevating intracellular amino acid pools that led to downstream amino-acid–dependent alterations in mitochondrial metabolism (Schuler *et al*, 2021; Raghuram & Hughes, 2024). Because glucose restriction slows translation and increases intracellular amino-acid pools, we initially hypothesized that MDC formation during glucose restriction might similarly arise from changes in amino-acid levels (Ashe *et al*, 2000; Schuler *et al*, 2021).

Consistent with this possibility, glucose restriction or carbon switching in synthetic medium, which contains substantially lower amino-acid concentrations than rich YPAD medium, failed to induce MDCs (Fig. 2A and Fig. S2A). Instead, mitochondria became highly fragmented (Fig. 2B). These observations initially suggested that amino-acid limitation might suppress MDC formation under these conditions. If this were the case, restoring amino-acid availability should rescue MDC formation. However, unlike previously reported MDC-inducing stresses such as rapamycin, concanamycin A (ConcA), and cycloheximide (CHX), addition of purified amino-acid sources such as casein or peptone failed to restore MDC formation. Only addition of yeast extract—which contains lipids, peptides, and metabolizable carbohydrates in addition to amino acids—readily restored MDC formation (Fig. 2C).

**Figure 2:**
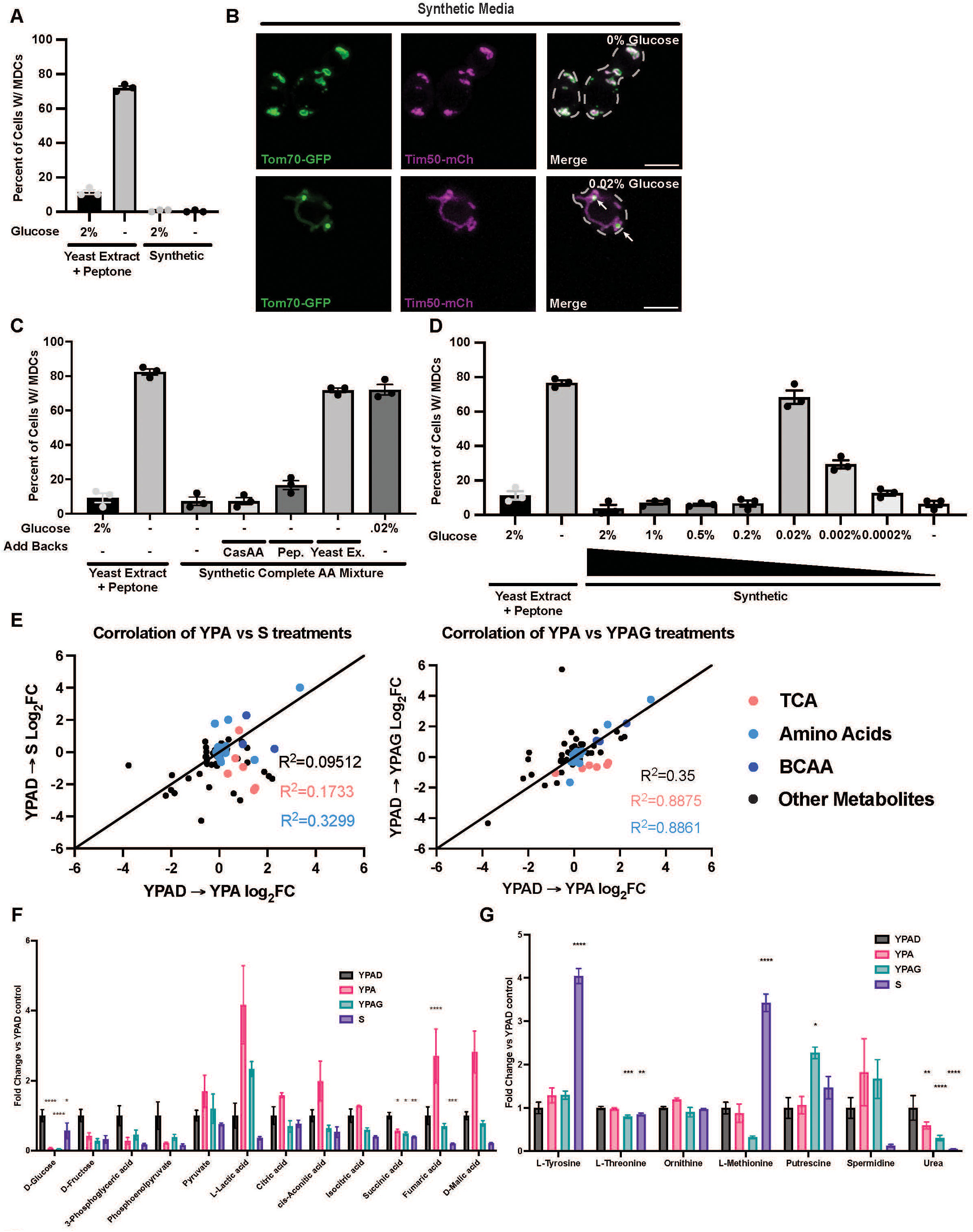
MDC formation during glucose restriction requires trace glucose but is independent of amino-acid availability. A. Quantification of the percentage of cells containing MDCs following glucose withdrawal in different media conditions. Cells were grown overnight in YPAD or synthetic (SD) media and shifted to the indicated media with or without glucose for 2 h. B. Super-resolution images of cells following glucose starvation (0% glucose, S) or glucose restriction (0.02% glucose, S + 0.02% glucose) for 2 h in synthetic media. Cells express Tom70-GFP and Tim50-mCherry. White arrows indicate MDCs. Scale bar, 5 μm. C. Quantification of MDC formation following 2 h add-back treatments. Cells were grown overnight in YPAD and shifted to the indicated conditions: YPAD (control), YPA (glucose restriction), S (no glucose), S + casamino acids (CasAA), S + peptone, S + yeast extract, or S + 0.02% glucose. D. Quantification of MDC formation across a glucose gradient. Cells were grown overnight in YPAD and shifted for 2 h to synthetic media containing the indicated glucose concentrations; YPAD and YPA serve as controls. E. Correlation analysis of metabolite log₂-fold changes following 2 h treatments. Left: glucose restriction (YPAD→YPA) versus glucose starvation (YPAD→S). Right: glucose restriction (YPAD→YPA) versus glycerol shift (YPAD→YPAG). Red points represent TCA cycle metabolites; blue, amino acid–related metabolites; dark blue, branched-chain amino acid–related metabolites; and black, other metabolites. Correlation coefficients (R^2^) are indicated. F. Analysis of central carbon metabolism metabolites following 2 h treatments (YPAD, YPA, YPAG, S). Values are shown as linear fold change relative to YPAD. n = 4; error bars represent mean ± SEM. G. Analysis of methionine and polyamine pathway metabolites under the same conditions as in (F). n = 4; error bars represent mean ± SEM. Quantification and statistics. Microscopy-based assays were performed with n = 3 independent experiments and 100 cells scored per replicate; error bars represent mean ± SEM. Statistical analysis for (F) and (G) was performed using two-way ANOVA with Holm–Šídák multiple-comparisons testing. False discovery rate (FDR) correction (q = 0.05) was applied.

Because purified amino-acid sources failed to rescue this response, but yeast extract did, we considered whether a non-amino-acid component of yeast extract, such as residual metabolizable carbon, was important. Previous studies have shown that synthetic medium with added glucose contains little to no residual glucose, whereas rich medium retains small amounts of metabolizable carbon after glucose removal due to components of yeast extract (Adachi *et al*, 2017; Ashe *et al*, 2000; Weber *et al*, 2020). We therefore asked whether it was the complete absence of glucose that was suppressing MDC formation in synthetic media. Consistent with this idea, adding trace amounts of glucose (0.02%) into glucose-free synthetic medium was sufficient to restore MDC formation and mitochondrial structure in this media type, producing MDC levels comparable to those observed during glucose restriction in rich medium (Fig. 2B–C). At glucose concentrations lower than 0.02%, mitochondria were fragmented and MDCs failed to form, indicating that MDC induction requires a minimal level of residual glucose present in the cell (Fig. 2D and Fig. S2B).

Consistent with the requirement for the presence of trace glucose for MDCs to form, MDC induction by other stimuli was likewise blocked under glucose-free conditions. For example, CHX-induced MDC formation was completely suppressed in glucose-free medium (Fig. S2C), suggesting that a minimal level of glucose is broadly required for MDC induction. This requirement does not appear to arise from the severe changes in mitochondrial morphology, as preventing mitochondrial fragmentation during complete glucose starvation by deleting the conserved mitochondrial fission GTPase Dnm1 failed to restore MDC formation under glucose-free conditions even though it prevented fragmentation (Fig. S2D-E). Although the basis for this glucose requirement remains unclear, similar observations have been reported for autophagy induction in yeast, where complete glucose starvation can prevent activation of the pathway in the presence of other starvation cues (Adachi *et al*, 2017). Together, these results indicate that MDC formation requires the presence of trace glucose across multiple induction conditions and suggest that MDC-permissive and MDC-nonpermissive glucose starvation conditions may represent fundamentally distinct metabolic states.

To better define how these carbon conditions differ metabolically, we performed untargeted GC-MS metabolomics on wild-type yeast cells grown overnight in YPAD and shifted for 2 h to control (YPAD), glucose restriction (YPA), carbon switch (YPAG), or synthetic glucose starvation (S). Global metabolomic responses were poorly correlated when comparing changes that occur upon a shift from YPAD to either YPA or S (R² ≈ 0.09), indicating that glucose restriction and complete glucose starvation produce distinct metabolic states (Fig. 2E). In contrast, shifting to YPA or YPAG showed moderate overall similarity in terms of metabolite changes (R^2^ ≈ 0.35), with substantially stronger concordance among amino acid and TCA cycle metabolites, indicating that these two MDC-permissive conditions share a common metabolic remodeling program (Fig. 2E).

At the pathway level, glycolytic and TCA cycle metabolites were broadly depleted in S relative to YPA or YPAG, whereas the MDC-permissive conditions maintained these pathways to a greater extent (Fig. 2F). Amino acid levels increased across multiple treatments, consistent with altered amino acid utilization and/or reduced anabolic demand, but this response alone was not sufficient to explain MDC formation (Fig. S2F). Instead, MDC-permissive conditions were associated with preservation of a partially active TCA-linked metabolic state, whereas complete glucose starvation produced broader suppression of biosynthetic metabolism (Fig. 2G).

Finally, we systematically examined the influence of amino-acid availability on glucose-restriction MDCs. MDCs were readily induced across rich (YPAD), synthetic, and minimal media conditions when 0.02% glucose was provided, and supplementation of casamino acids to match amino-acid levels found in rich medium did not enhance MDC formation (Fig. S2G). Moreover, MDCs were induced even during nitrogen starvation when trace glucose was present (Fig. S2H), demonstrating that amino-acid and nitrogen availability are not required for MDC formation under glucose restriction.

Together, these results show that glucose-restriction MDC formation requires reduced, but not absent, glucose availability and occurs under metabolic states that preserve aspects of central carbon metabolism. Unlike previously characterized MDC-inducing conditions such as concanamycin A, rapamycin, and cycloheximide, this response is independent of amino-acid availability, indicating that glucose restriction induces MDCs through a metabolically distinct remodeling state rather than through amino-acid accumulation alone.

### Snf1 is required for glucose restriction– and salt stress-induced MDCs

To understand what drives MDC formation during glucose-dependent metabolic transitions, we next examined signaling pathways that couple changes in carbon availability to MDC induction. Because glucose restriction and carbon-source switching engage well-defined glucose-responsive regulatory networks, we focused on major carbon-responsive pathways (Kunkel *et al*, 2019; Kayikci & Nielsen, 2015). Among the major glucose-sensing pathways, deletion of *SNF1*, a central energy-stress kinase that promotes respiratory metabolism, completely blocked MDC formation induced by glucose restriction and carbon-source switching, whereas deletion of the PKA-pathway regulators *IRA2*, *GPA2*, or *GPR1* had no effect (Fig. 3A-B; Fig. S3A). Because Snf1 activates mitochondrial biogenesis through transcriptional derepression, we next tested additional regulators of this pathway (Kayikci & Nielsen, 2015; Coccetti *et al*, 2018; Martinez-Ortiz *et al*, 2019; Fig. 3C). Deletion of *HXK2*, a repressor of glucose derepressed transcription, reduced MDC induction, whereas deletion of *REG1* or *REG2*, phosphatases that inhibit Snf1, had a negligible impact on MDC formation, possibly due to redundancy (Fig. S3B). We also tested whether Snf1 is required for MDC formation under other biogenesis-inducing conditions. Loss of *SNF1* blocked MDC induction during both NaCl and KCl stress (Fig. 3D), indicating that Snf1 broadly regulates MDC formation in response to metabolic stresses that promote mitochondrial biogenesis.

**Figure 3:**
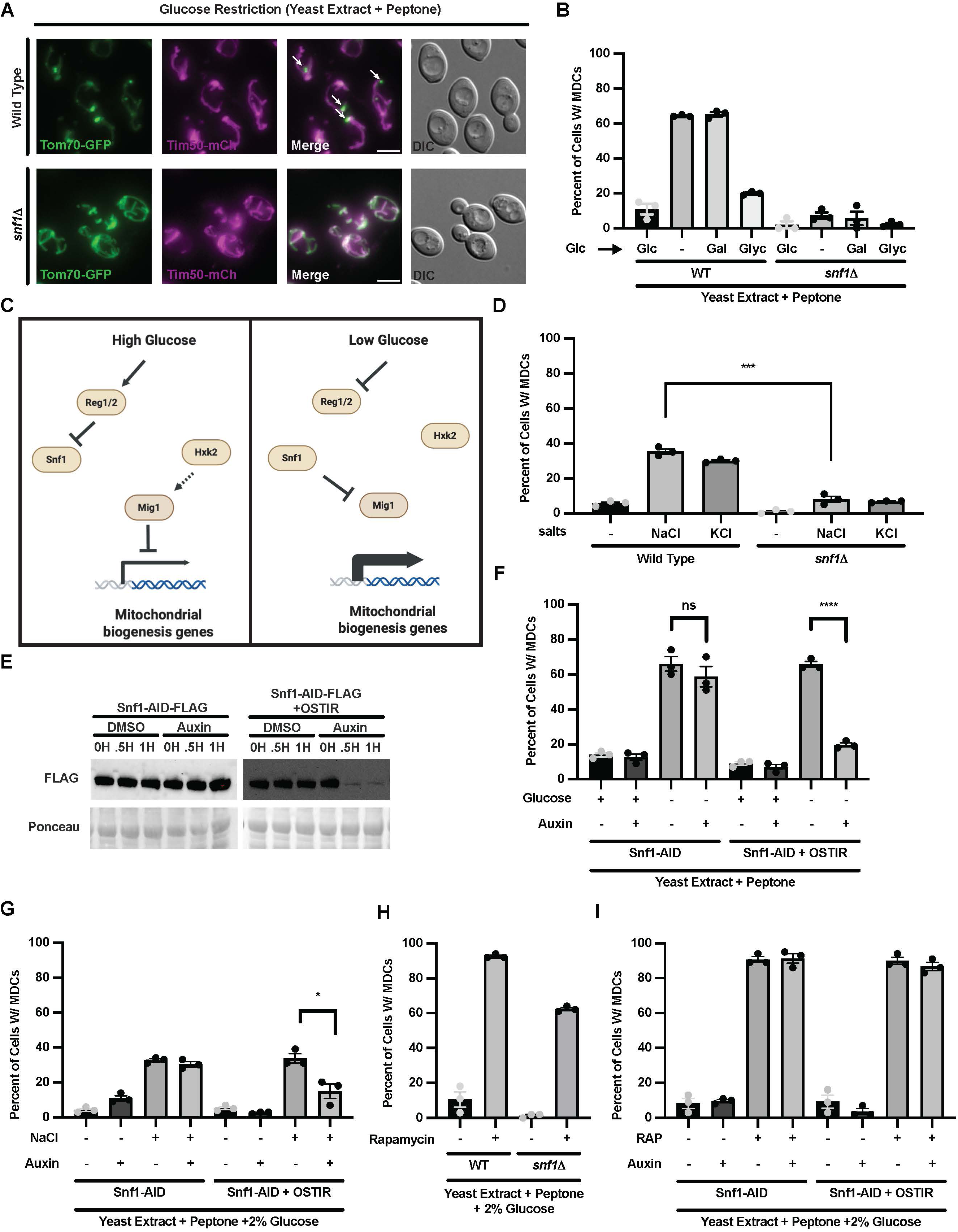
Snf1 is required for glucose restriction– and salt stress-induced MDCs. A. Widefield images of wild-type and *snf1Δ* cells following 2 h glucose restriction. Cells express Tom70-GFP and Tim50-mCherry. White arrows indicate MDCs. Scale bar, 5 μm. B. Quantification of MDC formation in wild-type and *snf1Δ* cells following 3 h carbon-source shifts. Cells were grown overnight in glucose (YPAD) and shifted to media containing glucose (Glc), no added carbon source (-), galactose (Gal), or glycerol (Glyc). C. Schematic of glucose sensing and repression through the Snf1 pathway. D. Quantification of MDC formation in wild-type and *snf1Δ* cells following 3 h salt stress (0.8 M NaCl or 1 M KCl). Wild-type and snf1Δ cells were analyzed in parallel. Wild-type data are reproduced from Figure 1G to facilitate direct comparison with the mutant. E. Immunoblot analysis of Snf1-AID-FLAG degradation following auxin treatment. Control (Snf1-AID-FLAG) and test (Snf1-AID-FLAG + His3-OsTIR1) strains were treated with DMSO or auxin for the indicated times. FLAG was detected by immunoblot; Ponceau staining serves as a loading control. F. Quantification of MDC formation following auxin-mediated Snf1 depletion. Cells were grown overnight in YPAD, pretreated with DMSO or auxin for 30 min, and subjected to glucose restriction for 2 h. G. Quantification of MDC formation in Snf1-AID strains following auxin-mediated Snf1 depletion. Cells were grown overnight in YPAD, pretreated with DMSO or auxin for 30 min, and subjected to salt stress for 3 h. H. Quantification of MDC formation in wild-type and *snf1Δ* cells following 2 h rapamycin treatment. I. Quantification of MDC formation in Snf1-AID strains following auxin-mediated Snf1 depletion. Cells were grown overnight in YPAD, pretreated with DMSO or auxin for 30 min, and subjected to rapamycin treatment for 2 h. Quantification and statistics. All MDC measurements (B, D, F-I) were performed with n = 3 independent experiments and 100 cells scored per replicate. Error bars represent mean ± SEM. Statistical significance was assessed using Welch’s two-tailed t-test.

Because MDCs form rapidly following glucose restriction, we asked whether Snf1 activity is acutely required for their induction. To test this, we used an auxin-inducible degron (AID) system to rapidly deplete Snf1 (Phanindhar & Mishra, 2023; Zahm *et al*, 2021). Auxin treatment for 30 minutes reduced Snf1-AID levels (Fig. 3E; Fig. S3C). Acute Snf1 depletion prior to treatment markedly impaired MDC induction during glucose restriction, salt stress, carbon switching, and 2-DG treatments whereas untreated Snf1-AID cells showed normal responses (Fig. 3F-G; Fig. S3D–F). These results demonstrate that Snf1 activity is acutely required for MDC formation under biogenesis-inducing conditions.

We next tested whether Snf1 dependence is specific to carbon and salt-stressed MDC induction. Neither deletion nor acute depletion of Snf1 affected MDC formation induced by rapamycin (Fig. 3H-I). Similarly, MDCs induced by concA or by constitutive overexpression of outer-membrane proteins (*SCM4* overexpression) were unaffected by Snf1 loss (Fig. S3G-H). Consistent with this, although *snf1Δ* did not impair *SCM4*-induced MDCs, it still blocked the additional increase in MDC formation triggered by glucose restriction in this scenario (Fig. S3I). Together, these findings indicate that Snf1 specifically regulates MDC formation driven by carbon and salt stress but is dispensable for MDCs induced by alterations in mTOR signaling, V-ATPase impairment, or direct increases in OMM protein load. These results identify Snf1 as a key regulator of carbon and salt stress-induced MDC formation and distinguish this pathway from Snf1-independent modes of MDC induction.

### Transcriptional derepression via Mig1 is required downstream of Snf1 in MDC induction

To identify the downstream effector(s) of Snf1 that drive MDC formation during glucose-dependent metabolic transitions, we examined Snf1-regulated pathways known to influence mitochondrial metabolism and remodeling. Snf1 has established roles in the regulation of autophagy, mitophagy, β-oxidation, lipid metabolism, and transcriptional control of glucose derepression (Usaite *et al*, 2009; Kayikci & Nielsen, 2015; Wei *et al*, 2016; Yao *et al*, 2020). Among these, only perturbation of the glucose derepression program consistently altered MDC induction during glucose restriction (Fig. 4A–C; Fig. S4A-B). Specifically, disruption of pathways downstream of Snf1 involved in β-oxidation (*FOX2*), stress and nutrient signaling (*RIM15*), autophagy (*ATG11*), or mitophagy (*ATG32*) had no detectable effect on glucose restriction–induced MDC formation (Fig. S4A). Likewise, perturbation of Snf1-regulated lipid metabolism did not phenocopy the specific requirement for Snf1 in glucose-responsive MDC induction. Acute repression of *ACC1*, a rate-limiting enzyme in de novo fatty acid synthesis and a well-established Snf1 target during glucose restriction (Wei *et al*, 2016), did not stimulate MDC formation and instead broadly inhibited MDC induction across multiple conditions (Fig. S4B-C, E). Similarly, treatment with cerulenin to impair de novo lipid synthesis reduced MDC formation across multiple inducers (Fig. S4D), indicating that lipid biosynthesis is generally required for MDC biogenesis but unlikely to represent the specific downstream pathway linking Snf1 signaling to glucose-responsive MDC induction. Together, these results identify transcriptional derepression as the primary downstream pathway coupling Snf1 activity to MDC formation.

**Figure 4:**
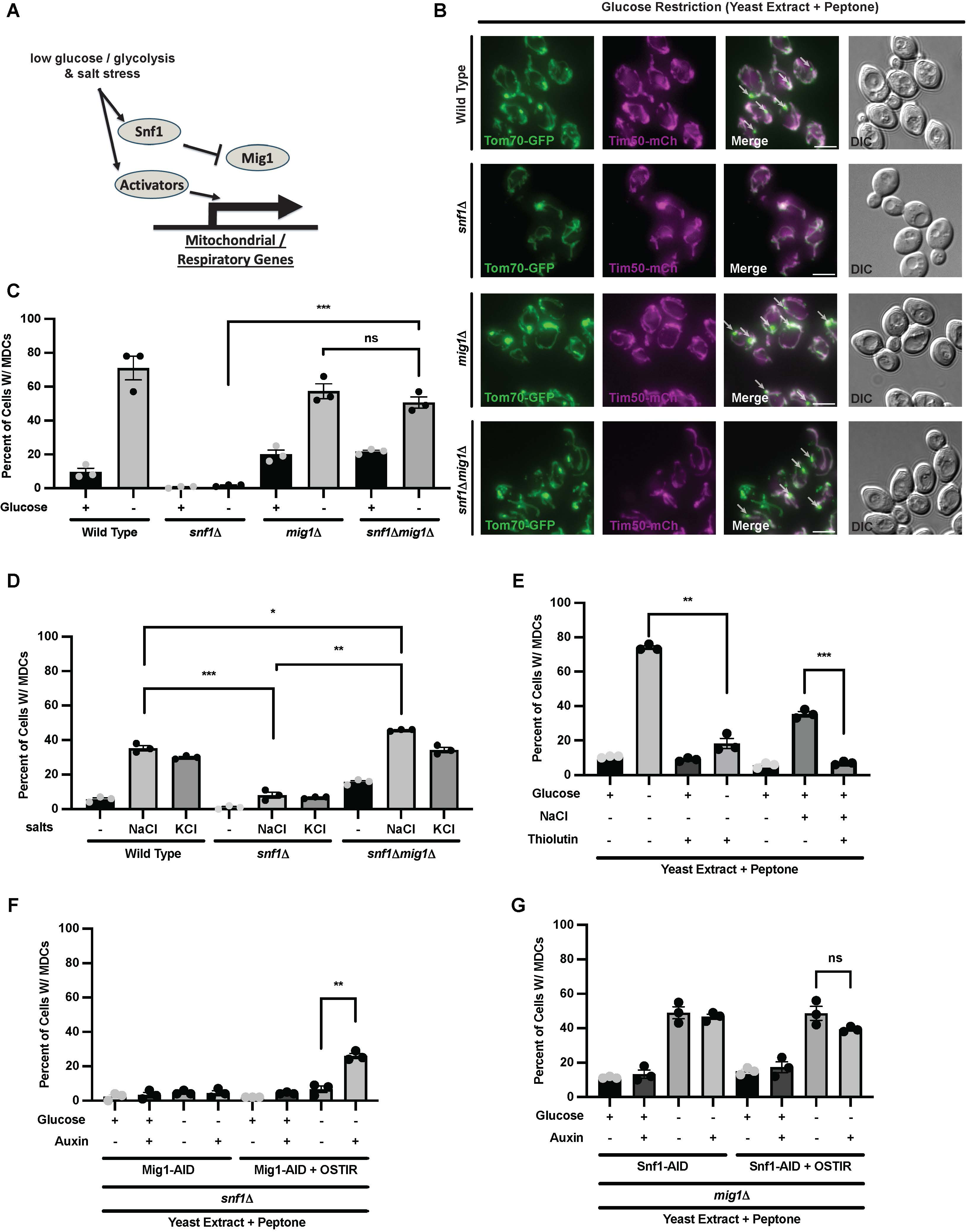
Mig1 derepression is required downstream of Snf1 for MDC induction. A. Schematic of the glucose repression pathway in yeast. B. Widefield images of wild-type, *snf1Δ*, *mig1Δ*, and *snf1Δ mig1Δ* cells following 2 h glucose restriction. Cells express Tom70-GFP and Tim50-mCherry. White arrows indicate MDCs. Scale bar, 5 μm. C. Quantification of MDC formation in wild-type, *snf1Δ*, and *snf1Δ mig1Δ* cells following 2 h glucose restriction. D. Quantification of MDC formation in wild-type, *snf1Δ*, and *snf1Δ mig1Δ* cells following 3 h salt stress (0.8 M NaCl or 1 M KCl). Wild-type and *snf1Δ* cells were analyzed in parallel with *snf1Δ mig1Δ* cells. The wild-type and *snf1Δ* data are reproduced from Figures 1G and 3D, respectively, to facilitate direct comparison. E. Quantification of MDC formation in wild-type cells following glucose restriction or NaCl stress in the presence or absence of thiolutin. Cells were treated for 2 h under glucose restriction conditions or for 3 h under NaCl stress. F. Quantification of MDC formation in *snf1Δ* MIG1-AID-FLAG cells with or without integrated HIS3::GPD-OsTIR1. Cells were grown overnight in YPAD, pretreated with DMSO or auxin for 30 min, and subjected to glucose restriction for 2 h. G. Quantification of MDC formation in *mig1Δ* SNF1-AID-FLAG cells with or without integrated HIS3::GPD-OsTIR1. Cells were grown overnight in YPAD, pretreated with DMSO or auxin for 30 min, and subjected to glucose restriction for 2 h. Quantification and statistics. All MDC measurements were performed with n = 3 independent experiments and 100 cells scored per replicate. Error bars represent mean ± SEM. Statistical significance was assessed using Welch’s two-tailed t-test.

Under glucose-rich conditions, the transcriptional repressor Mig1 contributes to repression of glucose-derepressed genes, including genes involved in alternative carbon metabolism and respiratory growth. Upon glucose limitation, Snf1 phosphorylates Mig1, promoting transcriptional derepression (Fig. 4A) (Treitel *et al*, 1998; Kayikci & Nielsen, 2015). Consistent with this model, deletion of *MIG1* fully restored MDC formation in *snf1Δ* cells under both glucose restriction and salt stress (Fig. 4B–D), demonstrating that relief of Mig1-mediated repression is sufficient to bypass the requirement for Snf1. Additionally, Snf1-dependent phosphorylation sites on Mig1 have been shown to be required for proper transcriptional derepression (Treitel *et al*, 1998). Consistent with this, CRISPR-generated Mig1 phospho-null mutants reduced glucose restriction-induced MDC formation, albeit to varying degrees (Fig. S4F).

To test whether active transcription is required for MDC formation, we inhibited transcription initiation using thiolutin (Pelechano & Pérez-Ortín, 2008). Thiolutin treatment prevented increased expression of mitochondrial proteins during glucose restriction and salt stress, and abolished MDC formation under both conditions (Fig. 4E; Fig. S4G). Because transcriptional inhibition has broad effects, we interpret this experiment conservatively. Nevertheless, acute thiolutin treatment during glucose restriction inhibited MDC formation within the same treatment window in which the Snf1–Mig1 axis is required, consistent with an acute requirement for ongoing transcription during MDC biogenesis (Pelechano & Pérez-Ortín, 2008; Mołoń *et al*, 2026).

Because MDC induction occurs rapidly following metabolic stress, we next asked whether Mig1 derepression is acutely required for MDC formation. To address this, we generated a Mig1-AID strain in a *snf1Δ* background and selectively degraded Mig1 using auxin (Fig. S4E). Acute Mig1 depletion alone was insufficient to induce MDCs; however, under glucose restriction, Mig1-AID degradation partially restored MDC formation in *snf1Δ* cells (Fig. 4F). We next performed the reciprocal experiment by degrading Snf1 in a *mig1Δ* background using a Snf1-AID allele (Fig. S4E). As expected, *mig1Δ* cells displayed MDC induction comparable to wild type under glucose restriction. Importantly, acute degradation of Snf1 in this background did not reduce MDC formation (Fig. 4G), phenocopying the *snf1Δ mig1Δ* double mutant. Together, these findings establish that MDC formation during glycolytic stress requires acute relief of Mig1-mediated transcriptional repression downstream of Snf1 signaling, identifying transcriptional derepression as the primary pathway coupling glucose-responsive metabolic remodeling to MDC formation.

### Transient induction of mitochondrial-targeted protein expression is a common theme among MDC inducers and sufficient to stimulate MDC formation

Because transcriptional derepression is required for MDC induction, and we have previously found that elevated expression of OMM proteins drives MDC formation, we next asked whether increased mitochondrial protein expression is a driver of MDC formation during metabolic transitions that stimulate MDCs. Acute upregulation of mitochondrial proteins occurs during both glucose restriction and salt stress (Fig. 1A, F), and this response is dependent on the Snf1–Mig1 regulatory axis (Fig. 5A). Importantly, we found that multiple previously characterized MDC-inducing conditions also triggered increased mitochondrial protein expression. Treatment with rapamycin or concA led to elevated levels of the outer-membrane proteins Om45 and Por1 with kinetics similar to MDC induction (Fig. S5A). In contrast to glucose restriction, rapamycin-induced mitochondrial protein expression was largely independent of Snf1, as protein levels were not substantially reduced in *snf1Δ* or Snf1-AID strains (Fig. S5B), consistent with the observation that rapamycin-induced MDC formation also does not require Snf1 signaling (Fig. 3H-I). Together, these results indicate that transient induction of mitochondrial-targeted protein expression is a shared feature of multiple MDC-inducing conditions, supporting a broader model in which elevated mitochondrial membrane protein load may represent a common factor in MDC formation.

**Figure 5:**
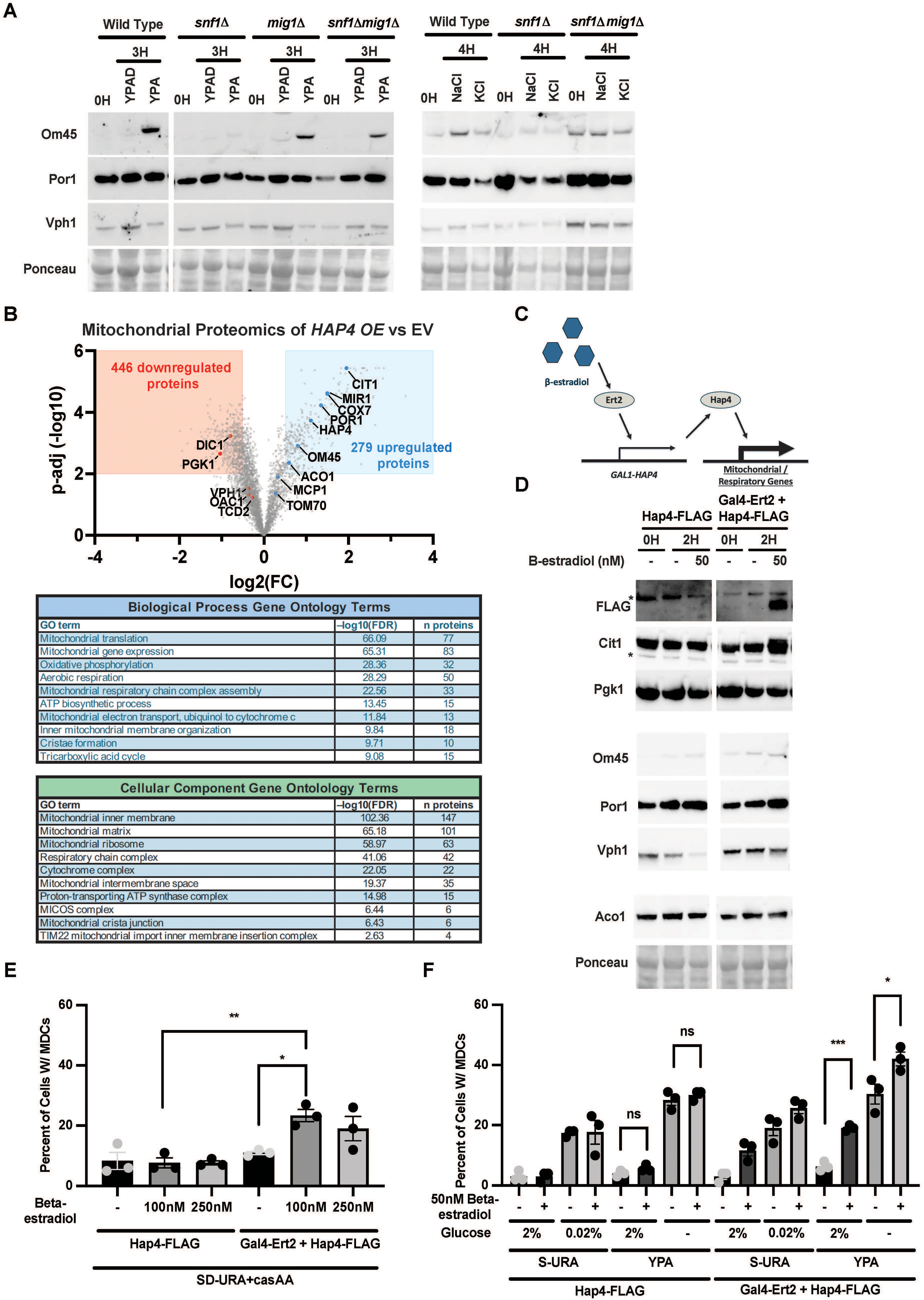
Transient induction of mitochondrial-targeted protein expression is a common theme among MDC inducers and sufficient to stimulate MDC formation. A. Immunoblot analysis of mitochondrial protein levels in wild-type, *snf1Δ*, *mig1Δ*, and *snf1Δ mig1Δ* cells following glucose restriction (3 h) or salt stress (0.8 M NaCl or 1 M KCl, 4 h). Om45 and Por1 are shown; Vph1 and Ponceau staining serve as loading controls. B. Mitochondrial proteomics analysis of Hap4 overexpression (OE) versus empty vector (EV). Top: volcano plot showing differentially regulated proteins using cutoffs of ≥1.5-fold change and FDR-adjusted p ≤ 0.01. Bottom: Gene Ontology (GO) enrichment analysis of upregulated proteins. C. Schematic of the β-estradiol–inducible Hap4 expression system. D. Immunoblot analysis of Hap4 induction following â-estradiol treatment. Strains expressing GAL1-HAP4-FLAG or GAL1-HAP4-FLAG + GAL4-ERT2 were treated with DMSO or â-estradiol as indicated. FLAG, Cit1, Om45, Por1, and Aco1 were detected; Pgk1, Vph1, and Ponceau staining serve as loading controls. The asterisk indicates a nonspecific band. E. Quantification of MDC formation in Hap4 expression strains following 3.5 h β-estradiol treatment at the indicated concentrations. Cells were maintained in SD-URA supplemented with casamino acids. F. Quantification of MDC formation in Hap4 expression strains under glucose-replete and glucose-restricted conditions with β-estradiol treatment. Cells were grown overnight in SD-URA and shifted to SD-URA, S-URA + 0.02% glucose, YPAD, or YPA as indicated for 3.5 h. Quantification and statistics. All MDC measurements were performed with n = 3 independent experiments and 100 cells scored per replicate. Error bars represent mean ± SEM. Statistical significance was assessed using Welch’s two-tailed t-test.

Because transcriptional derepression is required for MDC induction, and because multiple MDC-inducing conditions increase mitochondrial protein expression, we next asked whether acute activation of mitochondrial biogenesis is sufficient to trigger MDC formation in the absence of metabolic stress. Hap4 is a central activator of mitochondrial biogenesis and is sufficient to drive increased mitochondrial protein expression (Raghuram & Hughes, 2024; Lascaris *et al*, 2004). Indeed, proteomic analysis of isolated mitochondria from cells constitutively overexpressing *HAP4* revealed broad increases in mitochondrial proteins (Fig. 5B). To directly test acute sufficiency, we established a β-estradiol–inducible *HAP4* expression system (Costa *et al*, 2018) (Fig. 5C). Acute induction of *HAP4* increased levels of mitochondrial proteins, including Cit1, Aco1, Om45, and Por1 (Fig. 5D). Although modest increases were observed in untreated controls, induction upon *HAP4* activation was robust. Importantly, transient *HAP4* induction was sufficient to stimulate MDC formation, reaching ∼25% of cells after 3.5 hours (Fig. 5E). Although this response was lower than MDC induction during glucose restriction in rich medium, it was comparable to MDC levels observed under trace-glucose conditions in the same synthetic medium background (Fig. 5F). Moreover, glucose restriction enhanced MDC formation in combination with transient *HAP4* induction (Fig. 5F), suggesting that elevated mitochondrial protein expression cooperates with transient alterations in mitochondrial metabolic state to promote robust MDC biogenesis. In contrast, prolonged *HAP4* induction via overnight β-estradiol treatment suppressed MDC formation, phenocopying previously observed effects of chronic mitochondrial biogenesis and blocking MDC induction during rapamycin treatment (Fig. S5C; Raghuram & Hughes, 2024). Together, these results indicate that acute increases in mitochondrial protein synthesis are sufficient to stimulate MDC formation, that this response is enhanced when mitochondrial function is simultaneously perturbed, and that sustained or chronic biogenesis uncouples protein load from MDC induction. These findings support a model in which mitochondrial protein load and the metabolic adaptation state of the organelle together determine MDC sensitivity.

### MDC formation requires proper mitochondrial protein targeting

Having shown that acute mitochondrial protein synthesis can drive MDC formation, particularly when combined with metabolic perturbation, we next asked whether delivery of these proteins to mitochondria is required for MDC induction. Our previous observations indicated that disruption of mitochondrial import causes mislocalization of outer-membrane proteins such as Om45 to the endoplasmic reticulum, providing a context to test whether proper delivery of proteins to the OMM is critical for MDC formation (Shakya *et al*, 2021; Xiao *et al*, 2021). We therefore tested whether impairing the TOM complex receptors Tom70 and Tom71, which mediate targeting of hydrophobic mitochondrial proteins and are required for proper localization of multiple MDC cargoes, affects MDC induction during glucose restriction (Wiedemann & Pfanner, 2017; Schuler *et al*, 2021; Xiao *et al*, 2021). Deletion of both *TOM70* and *TOM71* abolished MDC formation observed with known MDC substrates Tcd2–GFP and Cox7–GFP under glucose restriction (Fig. 6A-B). In contrast, deletion of *TOM70* alone produced an intermediate phenotype, consistent with functional redundancy between Tom70 and Tom71. Correspondingly, mitochondrial proteins including Oac1-GFP and Om45-GFP were mislocalized in *tom70Δ tom71Δ* cells (Fig. 6C-F). Importantly, mitochondrial biogenesis-associated protein induction remained intact in *tom70Δ tom71Δ* strains during glucose restriction (Fig. 6G), indicating that loss of MDC formation does not arise from failure to activate the upstream glucose-responsive biogenic program. Together, these observations suggest that proper targeting of mitochondrial proteins to mitochondria is required for MDC formation, consistent with a model in which acute outer-membrane protein delivery drives MDC biogenesis during mitochondrial adaptation.

**Figure 6:**
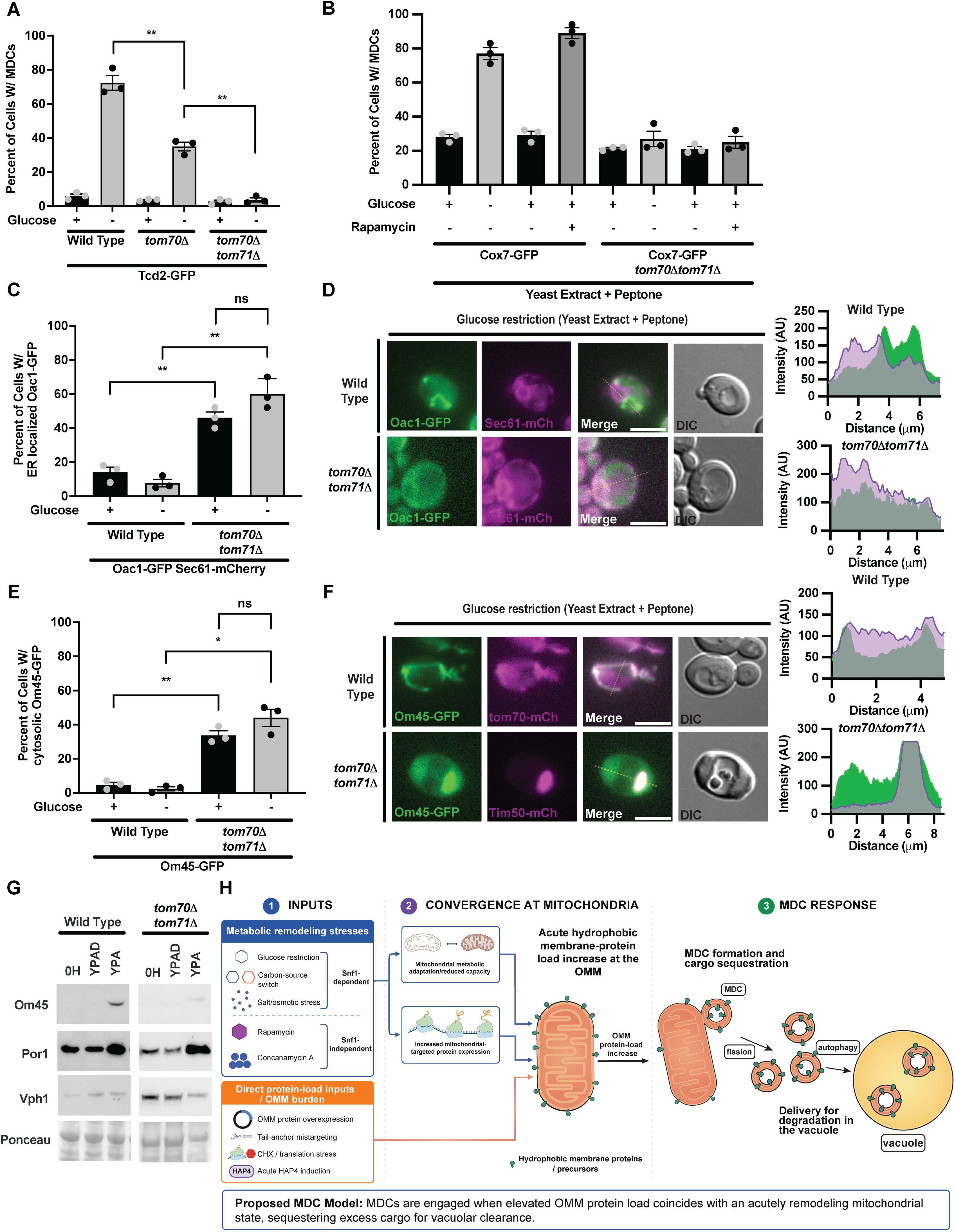
MDC formation requires Tom70/Tom71-dependent mitochondrial protein targeting. A. Quantification of MDC formation in wild-type, *tom70Δ*, and *tom70Δ tom71Δ* cells following 2 h glucose restriction. Cells express Tcd2-GFP and Tim50-mCherry. B. Quantification of MDC formation in wild-type and *tom70Δ tom71Δ* cells following 2 h glucose restriction or rapamycin treatment. Cells express Cox7-GFP and Tim50-mCherry. C. Quantification of the percentage of cells exhibiting ER-localized Oac1-GFP, as shown in (D), determined by colocalization with Sec61-mCherry. Cells were treated for 3 h in YPAD (control) or YPA (glucose restriction). D. Widefield images of Oac1-GFP localization in wild-type and *tom70Δ tom71Δ* cells following 3 h glucose restriction. Cells express Oac1-GFP and Sec61-mCherry. Line-scan intensity profiles corresponding to the yellow dashed lines are shown. Scale bar, 5 μm. E. Quantification of the percentage of cells exhibiting cytosolic Om45–GFP, as shown in (F), defined by diffuse cytosolic GFP signal. Cells were treated for 3 h in YPAD (control) or YPA (glucose restriction). F. Widefield images of Om45-GFP localization in wild-type and *tom70Δ tom71Δ* cells following 3 h glucose restriction. Cells express Om45–GFP with Tom70-mCherry or Tim50-mCherry. Line-scan intensity profiles corresponding to the yellow dashed lines are shown. Scale bar, 5 μm. G. Immunoblot analysis of mitochondrial protein levels in wild-type and *tom70Δ tom71Δ* cells following 3 h growth in YPAD or YPA. Om45 and Por1 are shown; Vph1 and Ponceau staining serve as loading controls. H. Proposed unified model for MDC formation. Snf1-dependent metabolic remodeling stresses, Snf1-independent nutrient/vacuolar stresses, and direct protein-load inputs converge on increased hydrophobic membrane-protein load at the OMM during acute mitochondrial remodeling. Tom70/Tom71-dependent targeting promotes engagement of these cargos with mitochondria, triggering MDC formation and cargo sequestration when protein load exceeds the adaptive capacity of the organelle. MDC cargos are subsequently delivered to the vacuole for degradation. Quantification and statistics. All microscopy-based assays were performed with n = 3 independent experiments and 100 cells scored per replicate. Error bars represent mean ± SEM. Statistical significance was assessed using Welch’s two-tailed t-test.

## Discussion

Metabolic transitions require cells to rapidly increase mitochondrial protein synthesis and remodel organelle composition to meet changing energetic demands (Pfanner *et al*, 2019; Palikaras *et al*, 2015). While the transcriptional programs that drive mitochondrial biogenesis are well established, how cells maintain mitochondrial proteostasis during these acute transitions has remained unclear (Kayikci & Nielsen, 2015; Posas *et al*, 2000; Boos *et al*, 2019; Kim *et al*, 2024). Here, we show that mitochondrial-derived compartments (MDCs), previously identified multilamellar structures derived from the outer mitochondrial membrane, are engaged during acute mitochondrial adaptation. Glucose restriction, carbon-source switching, and osmotic stress each induce a Snf1-dependent transcriptional program that increases mitochondrial protein expression, and under these conditions MDCs form robustly. MDC formation requires ongoing transcription and Tom70/Tom71-dependent mitochondrial protein targeting but remains modest when mitochondrial biogenesis is induced in the absence of metabolic perturbation and is suppressed after prolonged adaptation. Together, these findings support a model in which MDCs are most strongly engaged when increased mitochondrial protein influx converges with a mitochondrial state undergoing acute metabolic remodeling.

Based on these findings, we propose that MDCs may function as transient outer-membrane buffering domains that sequester membrane proteins delivered during periods when protein influx may exceed the immediate capacity of mitochondrial membrane organization and assembly pathways. Consistent with this model, disruption of mitochondrial protein targeting causes mitochondrial proteins to mislocalize and prevents MDC formation despite continued induction of mitochondrial protein expression. Thus, MDCs are not simply a consequence of increased protein synthesis, but require delivery and engagement of newly synthesized proteins with the mitochondrial outer membrane (Xiao *et al*, 2024, 2021). By concentrating these proteins within a discrete OMM-derived domain, MDCs may temporarily limit their accumulation across the broader mitochondrial network until the organelle adapts or the cargos are delivered to the vacuole for degradation (Wilson *et al*, 2024a).

This framework provides a possible unifying interpretation for prior MDC-inducing conditions (Fig. 6H). Many previously described triggers—including pharmacological inhibition of mTOR, V-ATPase disruption, translational stress, direct mitochondrial membrane protein overexpression, alterations in membrane lipid composition, and protein mistargeting—either increase hydrophobic membrane protein burden at the organelle, alter mitochondrial metabolic state, or both (Schuler *et al*, 2021; Raghuram & Hughes, 2024; Xiao *et al*, 2024; Wong *et al*, 2025; Wilson *et al*, 2024a). Thus, MDC formation appears to be favored when two conditions converge: elevated hydrophobic protein burden at the OMM and a mitochondrial state undergoing acute remodeling or adaptation. Consistent with this model, acute induction of mitochondrial biogenesis was sufficient to stimulate MDC formation, but the response was less robust than during metabolic perturbation and was further enhanced by glucose restriction. In contrast, chronic mitochondrial biogenesis suppressed MDC induction, suggesting that once fully adapted, mitochondria can accommodate increased protein expression without engaging MDC formation. MDC sensitivity therefore appears to reflect the relationship between protein load and the capacity of the organelle to accommodate that load, rather than either factor alone.

This interpretation is further supported by prior rapamycin-based genetic screening, which identified translation, transcriptional regulation, and chromatin/histone modification as modifiers of MDC induction, consistent with gene-expression programs contributing to outer-membrane remodeling (Xiao *et al*, 2024). MDCs therefore emerge not as the output of a single signaling pathway or metabolite, but as a broader OMM proteostasis response engaged when acute protein influx outpaces or is mismatched with mitochondrial adaptation.

Importantly, our study establishes MDCs as a physiologically relevant response associated with acute mitochondrial biogenesis rather than a phenomenon restricted to pharmacological perturbations. Under glucose restriction and related metabolic transitions, MDCs form during endogenous carbon-responsive remodeling programs. This places MDC formation early in the adaptive response, before cells resort to broader mitochondrial turnover pathways such as mitophagy (Palikaras *et al*, 2015; Kayikci & Nielsen, 2015). MDCs may therefore provide an intermediate layer of quality control that remodels selected regions of the OMM without requiring elimination of the entire organelle.

Several key questions remain. First, the molecular trigger for MDC formation is unknown. Although our data support a role for increased outer-membrane protein load, how this state is sensed and converted into membrane remodeling remains unclear. Second, the machinery that drives MDC formation has not been fully elucidated. Whether MDCs arise through regulated membrane deformation, lipid redistribution, or dedicated scission mechanisms remains an important question. Third, the functional consequences of MDC formation during mitochondrial biogenesis remain difficult to assess. Current approaches to block MDC formation disrupt upstream processes such as protein import or membrane organization, making it challenging to isolate the specific contribution of MDCs to mitochondrial homeostasis or directly test whether they buffer acute protein load.

In summary, we propose that MDCs are adaptive outer-membrane remodeling domains that respond to hydrophobic membrane protein load during acute mitochondrial adaptation. Rather than being triggered by protein load or metabolic stress alone, MDC formation is most strongly engaged when elevated outer-membrane protein burden coincides with a mitochondrial state undergoing active remodeling. By linking mitochondrial protein influx to membrane remodeling, this work positions MDCs as a physiologically relevant proteostasis pathway that may sequester and manage excess outer-membrane proteins while mitochondrial adaptation is underway.

## Materials and Methods

### Yeast strains

All yeast strains are derivatives of *S. cerevisiae* S288C (BY) (Baker Brachmann *et al*, 1998) and are listed in Table S1. Deletion strains were generated by one-step PCR-mediated gene replacement using pRS series vectors (Baker Brachmann *et al*, 1998; Sikorski & Hieter, 1989) and oligonucleotide pairs listed in Table S2. Correct gene deletions were confirmed by colony PCR across the chromosomal insertion site. Strains expressing proteins with C-terminal fluorescent tags were generated by one-step PCR-mediated endogenous tagging using standard techniques and oligo pairs listed in Table S2. Plasmid templates for fluorescent tagging were from the pKT series of vectors (Sheff & Thorn, 2004). Correct integrations were confirmed by colony PCR and by verifying proper localization of the fluorophore by microscopy. For diploid mutant strains, haploid parent strains were transformed with the desired constructs and mated on YPAD plates overnight. Colonies were subsequently selected on SD–Met–Lys plates.

For auxin-inducible degradation, proteins were tagged using one-step PCR-mediated C-terminal tagging with pHyg-AID*-6FLAG and corresponding oligo pairs listed in Table S2. Following tagging, His3-GPD-OsTIR1 (PM76) was integrated at the HIS3 locus to complete the degradation system (Zahm *et al*, 2021). The His3-GPD-OsTIR1 fragment was prepared by PmeI digestion and transformed into yeast.

Mig1 phospho-null mutants were generated using a CRISPR/Cas9-mediated genome editing approach in *Saccharomyces cerevisiae* as previously described (DiCarlo *et al*, 2013; Ryan *et al*, 2014). Guide RNAs targeting *MIG1* were designed proximal to the desired phosphorylation sites, and donor DNA templates containing serine-to-alanine substitutions (S→A) with ∼50–70 bp homology arms were synthesized to enable homology-directed repair (Table S2). Guide RNA target sites and associated PAM sequences were selected and cloned into pM696 by annealing and ligating double-stranded oligonucleotides containing the desired guide sequences. Functional guide RNAs were identified based on growth sensitivity on SD–URA following transformation. To prevent repeated Cas9 cleavage, silent mutations were introduced within the PAM or guide RNA recognition sequence where necessary. Guide RNA and donor sequences (Twist Biosciences) were cloned into plasmid pM1525 using Golden Gate assembly, and yeast strains expressing integrated Cas9 from pM693 digested with StuI and PmeI were transformed with these constructs. Following selection, clonal isolates were screened by PCR and confirmed by Sanger sequencing. The following Mig1 phospho-null mutants were generated: S1A (S381A), S2A (S222A, S278A), S3A (S222A, S278A, S381A), and S4A (S222A, S278A, S311A, S381A). Validated haploid Mig1 phospho-null strains were then mated to a *mig1Δ* strain to generate diploid strains for MDC assays.

The β-estradiol–inducible *HAP4* expression system was generated by integrating the GEM expression cassette into a chromosome I integration site (nt: 199456-199457) (Hughes & Gottschling, 2012). The Gal1–β-estradiol–Msn4 fragment was amplified using primers 5221/5222 from PJW-1663 (Addgene #112037) and transformed into BY4741 (Costa *et al*, 2018). This strain was subsequently transformed with PJW-HAP4-FLAG-1666 and maintained on SD–URA medium prior to treatment.

### Plasmids

The plasmids generated and used in this study are listed in Table S3. Most plasmids used in this study were generated by inserting PCR-amplified genes of interest into pRS series vectors (Baker Brachmann *et al*, 1998; Sikorski & Hieter, 1989) using Gibson assembly (Gibson *et al*, 2009). Expression was driven either by endogenous promoters or by the GPD promoter. The β-estradiol–inducible HAP4 expression plasmid (PJW-HAP4-FLAG-1666) was constructed by Gibson assembly. A synthetic HAP4-FLAG fragment (Twist Bioscience), containing flanking homology to PJW-1666 and a C-terminal FLAG tag, was inserted into PJW-1666 linearized with EcoRI and AscI. The assembled plasmid was transformed into competent *E. coli*, isolated via miniprep, and validated by restriction digestion and sequencing.

### Yeast cell culture and growth assays

Yeast cells were grown at 30°C for 15-16 h to mid-log phase (OD600 = 0.2–0.8) prior to all experiments. Unless otherwise indicated, cells were cultured in YPAD medium (1% yeast extract, 2% peptone, 0.005% adenine, and 2% glucose). Alternatively, cells were cultured in synthetic defined (SD) medium containing 0.67% yeast nitrogen base without amino acids, 2% glucose, and supplemented nutrients (0.072 g/L each adenine, alanine, arginine, asparagine, aspartic acid, cysteine, glutamic acid, glutamine, glycine, histidine, myo-inositol, isoleucine, lysine, methionine, phenylalanine, proline, serine, threonine, tryptophan, tyrosine, uracil, and valine; 0.369 g/L leucine; 0.007 g/L para-aminobenzoic acid). For amino acid–free conditions (MinD), cells were grown in medium containing 0.67% yeast nitrogen base without amino acids and 2% glucose. Where indicated, casamino acids were added at the time of treatment to a final concentration of 2%. Drugs and chemicals were used at the following concentrations unless otherwise indicated: rapamycin (200 nM), concanamycin A (500 nM), cycloheximide (500 nM), cerulenin (10 µM), thiolutin (10 µg/mL), sodium chloride (0.8 M), potassium chloride (1 M), and indole-3-acetic acid (1 mM) (Table S3). 2-deoxyglucose and β-estradiol were added at the indicated concentrations for each assay.

### MDC assays

For glucose restriction or media-switch assays, overnight log-phase cultures were centrifuged, washed 2–3 times with sterile ddH_2_O, and resuspended in the indicated media (YPAD, SD, MinD, or YPA [YPAD lacking 2% glucose]) or media containing specified metabolites for 2–3 h unless otherwise indicated. Carbon sources included trace glucose (0.02%), fructose (2%), glycerol (3%), galactose (2%), sucrose (2%), and raffinose (2%). For drug-based assays, cultures were incubated with DMSO or the indicated drug for 2 h. For plasmid-containing strains, overnight cultures grown in selective SD medium were diluted to OD600 = 0.1–0.2 in YPAD and allowed to grow for at least 1 h prior to induction. For combined treatments, pretreatment drugs were added prior to media switching, and drugs were re-added immediately following the switch. Prior to imaging, cells were harvested by centrifugation, washed once, and resuspended in 100 mM HEPES containing 5% glucose. Cells were plated onto slides at low volume to form a monolayer. Optical z-sections of live cells were acquired using a Zeiss Axio Imager M2 or Zeiss LSM800 Airyscan microscope. MDCs were quantified at 2–3 h after treatment. Data represent mean ± SEM from three biological replicates (n = 100 cells per replicate). MDCs were defined as Tom70-positive, Tim50-negative structures enriched relative to the mitochondrial network. In colocalization assays, MDCs were identified as large Tom70-enriched spherical structures prior to assessing protein colocalization.

### Fluorescence microscopy

Fluorescence microscopy was performed as described (English *et al*, 2020). Optical z-sections were acquired using a Zeiss Axio Imager M2 with a Zeiss Axiocam 506 camera and 100X or 63X oil-immersion objectives (NA 1.4), or a Zeiss LSM800 Airyscan microscope. Images were acquired using ZEN software and processed in Fiji (Schindelin *et al*, 2012). Airyscan images were processed using the automated Airyscan algorithm in ZEN and further analyzed in Fiji. Fluorophores are indicated in figure legends. Channels were minimally adjusted for visualization, and line-scan analyses were performed on unadjusted images. MDC diameters were measured from fields of cells imaged on the Zeiss Axio Imager M2 with a 63X oil-immersion objective. Images were processed by maximum-intensity projection, and diameters were measured using the line measurement tool in Fiji.

### Time-lapse imaging

For time-lapse imaging, overnight cultures were harvested, resuspended in SD medium, and loaded into flow chambers as described (Fees *et al*, 2017). After 15 min of incubation, chambers were washed and medium was exchanged to trace-glucose medium (S + 0.02% glucose). Flow chambers were constructed using microscope slides and coverslips coated with concanavalin A (Sigma-Aldrich; L7647) and sealed with Parafilm. Chambers were further sealed with melted Vaseline. Imaging was performed on a Zeiss LSM880 Airyscan microscope equipped with a temperature-controlled chamber set to 30°C. Images were processed using ZEN and Fiji as described above.

### Whole-cell lysate preparation and immunoblotting

Cell lysates and immunoblotting methods were adapted from (Wong *et al*, 2025). Two OD600 units of cells were lysed in 1 mL ice-cold 0.2 M NaOH for 10 min on ice. Trichloroacetic acid (100 µL) was added, and samples were incubated on ice for 5 min. Proteins were pelleted by centrifugation (13,000 rpm, 5 min) and resuspended in 2X SDS sample buffer (0.12M Tris-HCl, pH 6.8, 19% Glycerol, 0.15mM Bromophenol Blue, 3.8% SDS, and 0.05% β-mercaptoethanol). Samples were neutralized with 1 M Tris base (pH 11) and heated at 50°C for 30 min. Proteins were resolved by SDS–PAGE (4–12% gels, Bio-Rad) and transferred to nitrocellulose membranes. Membranes were blocked in TBST + 5% milk and incubated with primary antibodies (Table S3), followed by HRP-conjugated secondary antibodies (1:5000). Signals were detected using enhanced chemiluminescence and imaged with a Bio-Rad ChemiDoc MP system.

### Extraction of whole-cell metabolites from yeast

Whole-cell metabolite extraction was done as previously noted (Raghuram & Hughes, 2024) with specific alterations. Wild-type yeast cells were grown overnight in YPAD to a density of 0.3–0.5 × 10⁷ cells/mL, collected by centrifugation, and washed once with water. Cell pellets were resuspended in YPAD, YPA, YPAG, or S medium to represent control, glucose restriction, glycerol carbon switch, or glucose starvation conditions, respectively. Cells were cultured in the indicated media for 2 h, collected by centrifugation for 3 min at 5,000 × g, washed once with water, and flash frozen in liquid nitrogen.

Whole cell metabolites were extracted from yeast cell pellets as previously described, with slight modifications (Canelas *et al*, 2009). Briefly, 0.4 µg of the internal standard succinic-d4 acid was added to each sample. Next, 1 mL of boiling 75% EtOH was added to each pellet, followed by vortex mixing and incubation at 90°C for 3 min with intermittent vortexing. Cell debris was removed by centrifugation at 7,000 × g for 5 min at −10°C. Supernatants were transferred to new tubes and dried en vacuo. Process blank samples were prepared using extraction solvent without a cell pellet.

### GC-MS analysis

Non-targeted metabolomics GC-MS analysis was done as previously described in (Raghuram & Hughes, 2024). GC-MS analysis was performed using an Agilent 5977B GC-MS MSD-HES equipped with an Agilent 7693A automatic liquid sampler. Dried samples were suspended in 40 µL of 40 mg/mL O-methoxylamine hydrochloride (MOX; MP Bio #155405) in dry pyridine (EMD Millipore #PX2012-7) and incubated for 1 h at 37°C in a sand bath. 25 µL of this solution was added to autosampler vials. For pooled quality-control (QC) samples, 10 µL of the remaining solution from each sample was combined, and 25 µL of the pooled QC was added to autosampler vials. Sixty microliters of N-methyl-N-trimethylsilyltrifluoracetamide (MSTFA with 1% TMCS; Thermo #TS48913) was added automatically via the autosampler and incubated for 30 min at 37°C. After incubation, samples were vortexed, and 1 µL of the prepared sample was injected into the gas chromatograph inlet in split mode with the inlet temperature held at 250°C. A 5:1 split ratio was used for analysis of the majority of metabolites. Metabolites that saturated the instrument at the 5:1 split ratio were analyzed at a 50:1 split ratio. The gas chromatograph had an initial temperature of 60°C for 1 min, followed by a 10°C/min ramp to 325°C and a 10 min hold. A 30 m Agilent Zorbax DB-5MS column with a 10 m Duraguard capillary column was used for chromatographic separation. Helium was used as the carrier gas at a flow rate of 1 mL/min.

Data were collected using MassHunter software (Agilent). Metabolites were identified, and peak areas were recorded using MassHunter Quant. These data were transferred to an Excel spreadsheet (Microsoft, Redmond, WA). Metabolite identity was established using a combination of an in-house metabolite library developed from purchased standards, the NIST library, and the Fiehn library. Values for each metabolite were normalized to the internal standard in each sample, normalized to total signal, and displayed as fold change relative to the control sample. Data were analyzed using the MetaboAnalyst software tool (Pang *et al*, 2021). Log₂ fold-change values from MetaboAnalyst were used for correlation analyses. Statistical analysis across metabolites was performed using two-way ANOVA with Holm–Šídák multiple-comparisons testing and verified using the two-stage linear step-up procedure of Benjamini, Krieger, and Yekutieli for FDR correction. All steady-state metabolite data and MetaboAnalyst input files are provided in Table S4.

### Isolation of yeast mitochondria

Isolation of yeast mitochondria was done as described in (Balasubramaniam *et al*, 2026). Yeast were grown overnight to log-phase (OD₆₀₀= 0.4–0.8). Cells were collected by centrifugation, washed with distilled water, and weighed. Cell pellets were resuspended in dithiothreitol (DTT) buffer (0.1 M Tris, 10 mM DTT, pH 9.4) at 2 mL per gram of pellet and incubated for 20 min at 30°C under constant rotation. After centrifugation, DTT-treated cells were washed once in 1.2 M sorbitol and resuspended in sorbitol phosphate buffer (6.7 mL per gram of pellet) containing 2 mg lyticase per gram of pellet. Cells were incubated for 30–45 min at 30°C with constant rotation to digest the cell wall and generate spheroplasts. Following lyticase treatment, spheroplasts were isolated by centrifugation and lysed by mechanical disruption in homogenization buffer (0.6 M sorbitol, 10 mM Tris, pH 7.4, 1 mM EDTA, pH 8.0 adjusted with KOH, 0.2% BSA, and 1 mM PMSF) at 13.3 mL per gram of pellet at 4°C. Cell debris was removed from the homogenate twice by centrifugation at 2,000 × g for 5 min at 4°C, and mitochondria were pelleted at 17,500 × g for 12 min at 4°C. The mitochondrial pellet was resuspended in SEM buffer (250 mM sucrose, 1 mM EDTA, pH 8.0 adjusted with KOH, and 10 mM 3-(N-morpholino) propanesulfonic acid [MOPS], pH 7.2), reisolated by differential centrifugation as described above, and subsequently resuspended in SEM buffer. Mitochondrial protein concentration was determined by bicinchoninic acid (BCA) assay. For proteomics analysis, 100 µg of mitochondria by protein mass was pelleted, flash frozen in liquid nitrogen, and stored at −80°C. Four biological replicates were analyzed per condition, and all samples were processed and acquired within the same LC-MS run.

### Mass spectrometry sample preparation

Samples for protein analysis were prepared as previously described (Navarrete-Perea *et al*, 2018; Li *et al*, 2021). Proteomes were extracted using a buffer containing 200 mM EPPS, pH 8.5, 8 M urea, 0.1% SDS, and protease inhibitors. Following lysis, 25 µg of each proteome was reduced with 5 mM TCEP. Cysteine residues were alkylated using 10 mM iodoacetamide for 20 min at room temperature in the dark. Excess iodoacetamide was quenched with 10 mM DTT. Buffer exchange was performed using a modified SP3 protocol (Hughes et al, 2019). Briefly, ∼250 µg of Cytiva SpeedBead Magnetic Carboxylate Modified Particles (65152105050250 and 4515210505250), mixed at a 1:1 ratio, was added to each sample. Ethanol was added to each sample to achieve a final concentration of at least 50%. Samples were incubated with gentle shaking for 15 min and washed three times with 80% ethanol. Protein was eluted from SP3 beads using 200 mM EPPS, pH 8.5, containing Lys-C (Wako, 129-02541). Samples were digested overnight at room temperature with vigorous shaking. The next morning, trypsin was added to each sample, and samples were incubated for an additional 6 h at 37°C. Acetonitrile was added to each sample to achieve a final concentration of ∼33%. Each sample was labeled in the presence of SP3 beads with ∼62.5 µg of TMTPro reagents (Thermo Fisher Scientific). Following confirmation of satisfactory labeling (>97%), excess TMT was quenched by addition of hydroxylamine to a final concentration of 0.3%. The full volume from each sample was pooled, and acetonitrile was removed by vacuum centrifugation for 1 h. The pooled sample was acidified, and peptides were desalted using a Sep-Pak 50 mg tC18 cartridge (Waters). Peptides were eluted in 70% acetonitrile, 1% formic acid, and dried by vacuum centrifugation.

### Basic pH reversed-phase separation (BPRP)

TMT-labeled peptides were solubilized in 5% acetonitrile/10 mM ammonium bicarbonate, pH 8.0, and ∼300 µg of TMT-labeled peptides was separated using an Agilent 300Extend-C18 column (3.5 µm particle size, 4.6 mm inner diameter, 250 mm length). An Agilent 1260 binary pump coupled with a photodiode array (PDA) detector (Thermo Scientific) was used to separate the peptides. A 45 min linear gradient from 10% to 40% acetonitrile in 10 mM ammonium bicarbonate, pH 8.0, at a flow rate of 0.6 mL/min separated the peptide mixture into 96 fractions collected every 36 s. The 96 fractions were consolidated into 24 samples in a checkerboard fashion and dried to completion by vacuum centrifugation. Each sample was desalted using StageTips and redissolved in 5% formic acid/5% acetonitrile for LC-MS3 analysis.

### Liquid chromatography and tandem mass spectrometry

Mass spectrometric data were collected on an Orbitrap Eclipse mass spectrometer coupled to a Proxeon NanoLC-1000 UHPLC (Thermo Fisher Scientific). The 100 µm capillary column was packed in-house with 35 cm of Accucore 150 resin (2.6 μm, 150Å; ThermoFisher Scientific). Data were acquired for 120 min per run. A FAIMS device was enabled during data collection and compensation voltages were set at –40V, –60V, and – 80V (Schweppe et al, 2019). MS1 scans were collected in the Orbitrap (resolution – 60,000; scan range – 400-1600 Th; automatic gain control (AGC) – 4×105, maximum ion injection time – automatic). MS2 scans were collected in the Orbitrap following higher-energy collision dissociation (HCD; resolution – 50,000; AGC – 1.25×105; normalized collision energy – 36; isolation window – 0.5 Th; maximum ion injection time – 86 ms.

### Mass spectrometry data analysis

Database searching included all entries from the Saccharomyces cerevisiae UniProt Database (downloaded in June 2023). The database was concatenated with one composed of all protein sequences for that database in the reversed order (Elias & Gygi, 2007). Raw files were converted to mzXML, and monoisotopic peaks were re-assigned using Monocle (Rad *et al*, 2021). Searches were performed with Comet (Eng *et al*, 2013) using a 50-ppm precursor ion tolerance and fragment bin tolerance of 0.02. TMTpro labels on lysine residues and peptide N-termini (+304.207 Da), as well as carbamidomethylation of cysteine residues (+57.021 Da) were set as static modifications, while oxidation of methionine residues (+15.995 Da) was set as a variable modification. Peptide-spectrum matches (PSMs) were adjusted to a 1% false discovery rate (FDR) using a linear discriminant, after which proteins were assembled further to a final protein-level FDR of 1% analysis (Huttlin *et al*, 2010). TMT reporter ion intensities were measured using a 0.003 Da window around the theoretical m/z for each reporter ion. Proteins were quantified by summing reporter ion counts across all matching PSMs. More specifically, reporter ion intensities were adjusted to correct for the isotopic impurities of the different TMTpro reagents according to manufacturer specifications. Peptides were filtered to exclude those with a summed signal-to-noise (SN) < 180 across all TMT channels and < 0.5 precursor isolation specificity. The signal-to-noise (S/N) measurements of peptides assigned to each protein were summed (for a given protein)._The mass spectrometry proteomics data have been deposited to the ProteomeXchange Consortium via the PRIDE (Perez-Riverol *et al*, 2025) partner repository with the dataset identifier PXD078768. Table S5 provides a summary of peptide counts and signal-to-noise (S/N) ratios across all identified proteins in all samples.

### Data visualization of mitochondrial proteomics

The EV and HAP4OE datasets were included for downstream analysis. Raw proteomics peak intensity data were analyzed using MetaboAnalyst. Initial data integrity checks revealed no errors, and data filtering was performed using default settings. Statistical preprocessing was conducted using the Statistical Analysis (one factor) module with no sample normalization, square root transformation, and Pareto scaling. Normalization quality was assessed by visual inspection of the resulting distribution plots, which demonstrated an approximately Gaussian distribution. Differential protein abundance between conditions was assessed using volcano plot analysis. Significance thresholds were set at ≥1.5-fold change and an adjusted p-value (FDR) ≤ 0.01. The resulting table of significantly altered proteins was exported from MetaboAnalyst and used for downstream visualization in GraphPad Prism. GO category analysis of HAP4OE versus EV mitochondrial proteomics data was performed using TMT Mosaic. Upregulated and downregulated protein sets were defined using cutoffs of ≥1.5-fold change and FDR-adjusted p ≤ 0.01, yielding 279 upregulated and 446 downregulated proteins. GO analysis of upregulated HAP4OE proteins was performed using the PANTHER Classification System statistical overrepresentation test (Mi *et al*, 2019). Upregulated and downregulated protein lists and GO-analysis outputs are provided in Table S6.

## Quantification and statistical analysis

The number of replicates, definition of n, and statistical tests are indicated in figure legends. Most experiments represent mean ± SEM from three biological replicates (n = 100 cells per replicate). Statistical analyses were performed using GraphPad Prism. All other statistical analyses are provided in the individual method sections above.

## Data availability

All reagents used in this study are available upon request. All other data reported in this paper will be shared by the lead contact upon request. This paper does not report original code. Any additional information required to reanalyze the data reported in this paper is available from the lead contact upon request.

## Acknowledgments

We thank members of the A.L. Hughes group for discussion and manuscript comments. We thank members of the Matthew P. Miller laboratory for providing reagents and support for the development of auxin-inducible degradation systems and CRISPR/Cas9 editing. Research was supported by NIH grant F31GM153135 to B.J.P., NIH grants GM119694 and AG061376 to A.L.H., and a Utah Graduate Research Fellowship to S.S.B. Metabolomics analysis was performed at the Metabolomics Core Facility at the University of Utah. Mass spectrometry equipment was obtained through NCRR Shared Instrumentation Grants 1S10OD016232-01, 1S10OD018210-01A1, and 1S10OD021505-01. We thank Quentin Pearce and James Cox for assistance with GC-MS instrument operation and metabolomics analysis. Proteomics analysis was conducted at the Thermo Fisher Scientific Center for Multiplexed Proteomics at Harvard Medical School. We thank Jonathan Van Vranken for performing the TMT-based proteomics experiments and Paul Stewart for assistance with subsequent data analysis and visualization. The content is solely the responsibility of the authors and does not necessarily represent the official views of the National Institutes of Health.

## Author Contributions

Conceptualization, B.J.P. and A.L.H.; methodology, B.J.P.; formal analysis, B.J.P.; investigation, B.J.P. and S.S.B.; writing – original draft, B.J.P.; writing – review and editing, B.J.P. and A.L.H.; visualization, B.J.P. S.S.B. and A.L.H.; supervision, A.L.H.; funding acquisition, B.J.P., S.S.B., and A.L.H.

## Online Supplemental Material

Movie S1 shows a 1 h time-lapse of wild-type cells during glucose restriction in YPA. Tom70–GFP is shown in green and Tim50–mCherry is shown in magenta. Table S2 lists the oligonucleotide sequences used in this study. Table S3 lists the plasmids, reagents, supplies, and software used in this study. Table S4 contains the raw metabolomics data and MetaboAnalyst input files. Table S5 contains the TMT-based quantitative proteomics dataset, including peptide counts and scaled signal-to-noise values for all identified proteins across all samples. Table S6 contains the GO analysis of proteins upregulated and downregulated in HAP4-overexpressing cells compared with empty-vector controls.

## Supplemental Figure Legends

**Figure S1:**
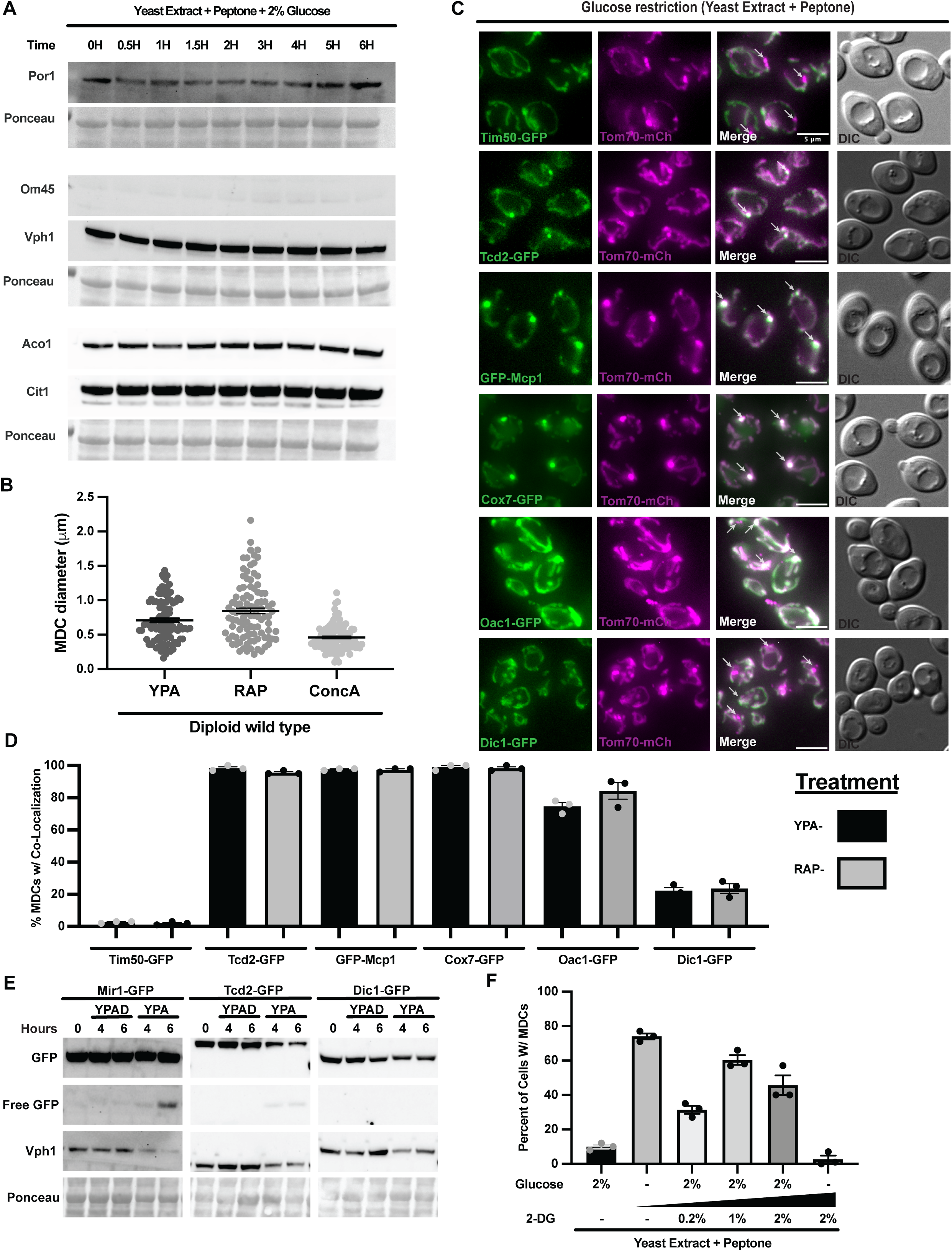
Characterization of MDCs induced by mitochondrial biogenesis–activating conditions (related to Figure 1) A. Immunoblot time course of mitochondrial protein levels in wild-type cells grown in glucose-replete conditions (YPAD) over 0-6 h. Por1, Om45, Aco1, and Cit1 are shown; Vph1 and Ponceau staining serve as loading controls. B. Quantification of MDC diameter under different induction conditions. Wild-type diploid cells were treated with glucose restriction (YPA), rapamycin (RAP), or concanamycin A (ConcA). C. Widefield images of MDC cargo colocalization following 2 h glucose restriction. Cells express Tom70–mCherry to label MDCs and the indicated GFP-tagged proteins (Tim50, Tcd2, Mcp1, Cox7, Oac1, or Dic1). White arrows indicate MDCs. Scale bar, 5 μm. D. Quantification of MDC cargo colocalization, as shown in (C). Colocalization was scored when MDCs overlapped with the indicated GFP signal. E. Immunoblot analysis of GFP processing for MDC cargo proteins following glucose restriction for the indicated times. Strains expressing Mir1-GFP, Tcd2-GFP, or Dic1-GFP were analyzed. Free GFP indicates proteolytic cleavage; GFP was detected by immunoblot, and Vph1 and Ponceau staining serve as loading controls. F. Quantification of MDC formation following 2 h treatment with 2-deoxy-D-glucose (2-DG) at the indicated concentrations. Quantification and statistics. For MDC frequency measurements, n = 3 independent experiments with 100 cells scored per replicate; for diameter measurements (B), n = 100 MDCs per condition. Error bars represent mean ± SEM.

**Figure S2:**
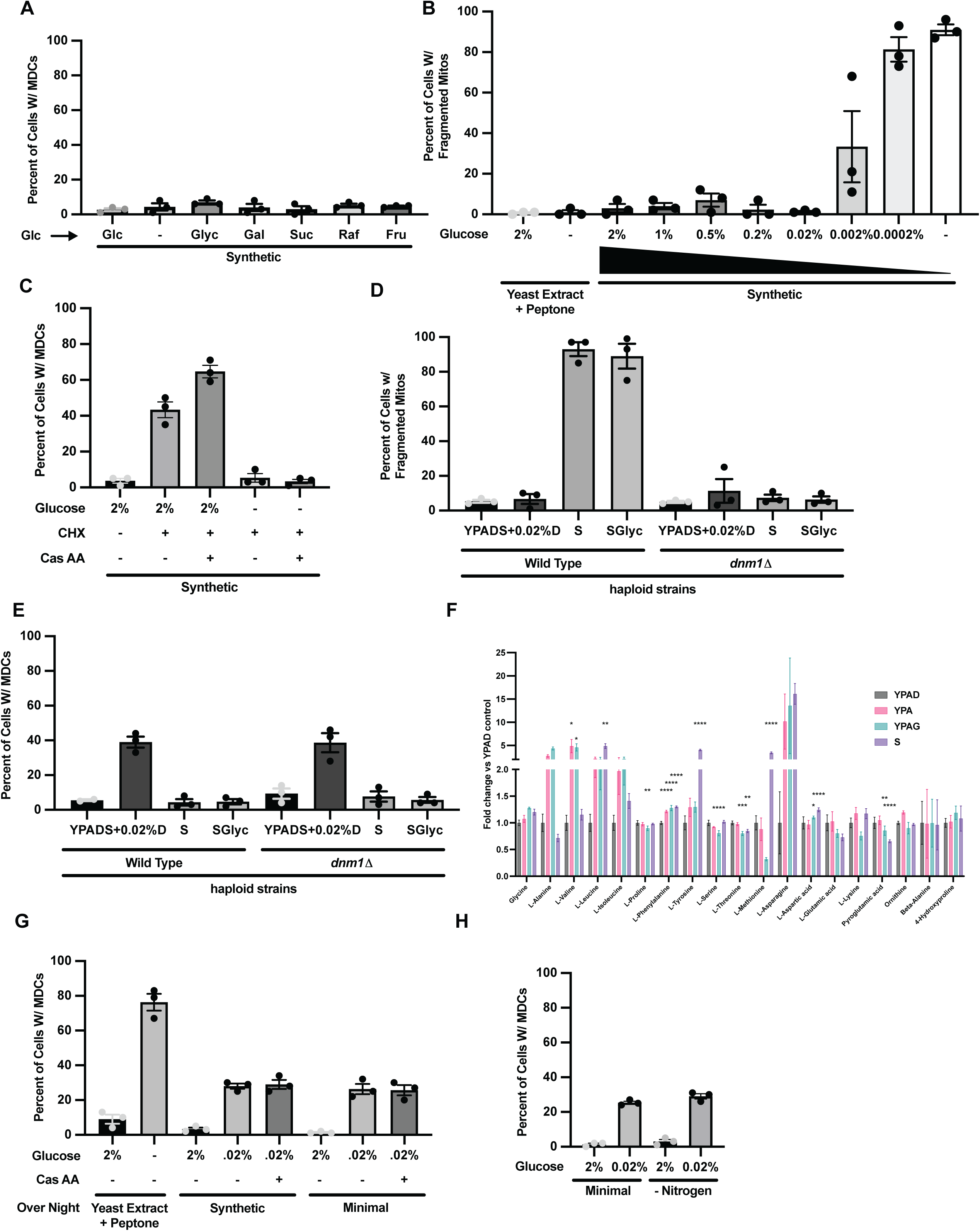
Further characterization of the trace-glucose requirement and amino acid independence of MDC formation (related to Figure 2) A. Quantification of MDC formation following 3 h carbon-source shifts in synthetic media. Cells were grown overnight in synthetic media with 2% glucose and shifted to media containing glucose (Glc), no added carbon source (-), glycerol (Glyc), galactose (Gal), sucrose (Suc), raffinose (Raf), or fructose (Fru). B. Quantification of the percentage of cells exhibiting fragmented mitochondria (see 2B and 2D). C. Quantification of MDC formation following 2 h cycloheximide (CHX) treatment under varying glucose and amino acid conditions. Cells were grown overnight in synthetic media and shifted to the indicated conditions. D. Quantification of mitochondrial fragmentation in wild-type and *dnm1Δ* cells under different media conditions. Cells were grown overnight in YPAD and shifted for 2 h to YPAD, synthetic media with 0.02% glucose (S + 0.02% glucose), synthetic media without carbon (S), or synthetic media with glycerol (S + glycerol). E. Quantification of MDC formation in wild-type and *dnm1Δ* cells under the conditions described in (D). F. Analysis of whole-cell amino acid metabolites following 2 h treatments (YPAD, YPA, YPAG, S), shown as linear fold change relative to YPAD. G. Quantification of MDC formation under trace-glucose conditions with or without amino acid supplementation. Prototrophic cells were grown overnight in YPAD, synthetic (SD), or minimal media and shifted for 2 h to the indicated conditions, including glucose restriction (0.02% glucose) and casamino acid supplementation. H. Quantification of MDC formation under trace-glucose conditions in minimal or nitrogen-starvation media. Cells were grown overnight in minimal media with 2% glucose and shifted for 2 h to the indicated conditions. Quantification and statistics. Microscopy-based assays (A–E, G–H) were performed with n = 3 independent experiments and 100 cells scored per replicate. Error bars represent mean ± SEM. Statistical analysis for (F) was performed using two-way ANOVA with Holm–Šídák multiple-comparisons testing; false discovery rate (FDR) correction (q = 0.05) was applied.

**Figure S3:**
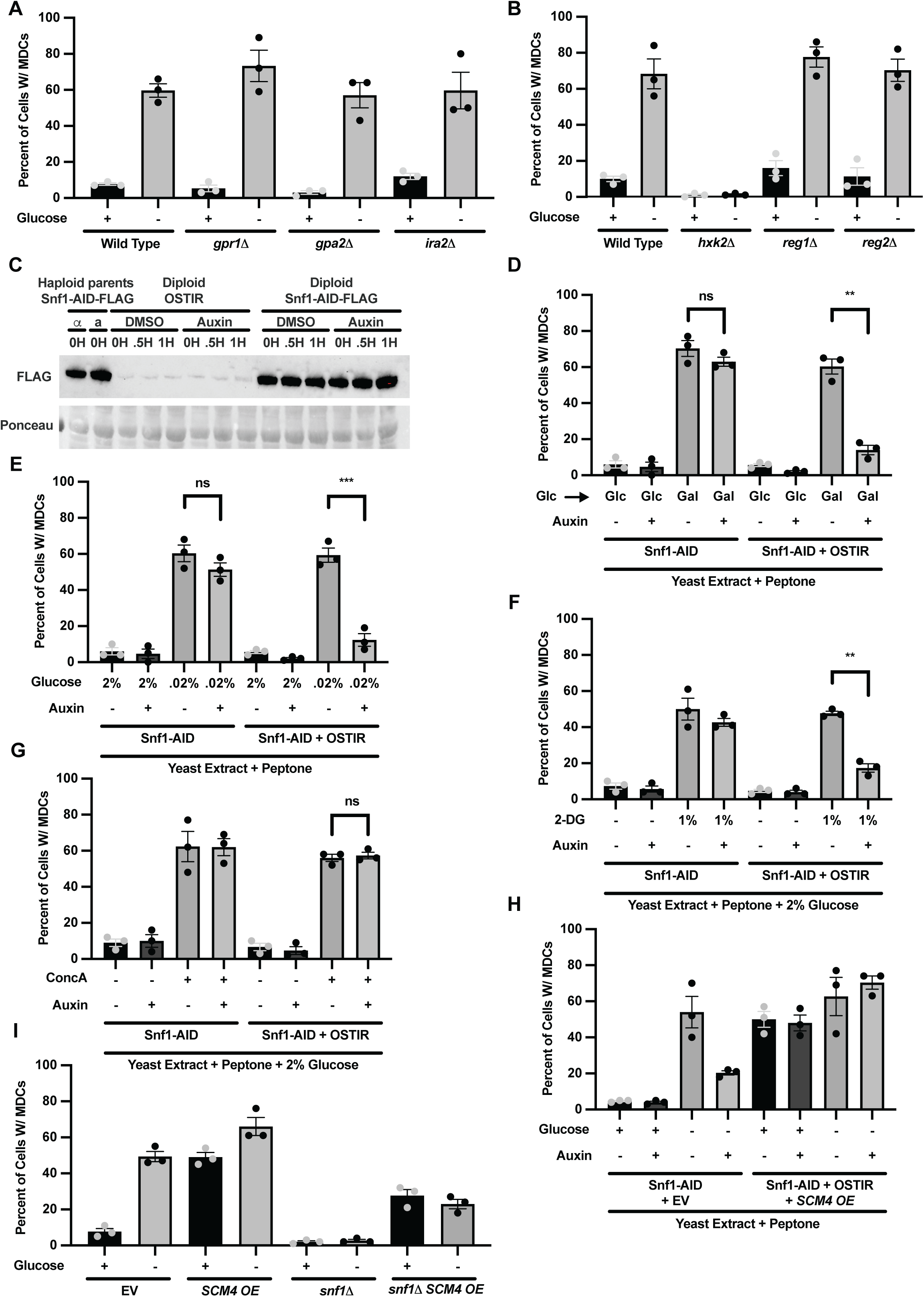
Snf1 is specifically and acutely required for MDC formation induced by mitochondrial biogenesis–activating conditions (related to Figure 3) A. Quantification of MDC formation in PKA-pathway mutants following 2 h glucose restriction. B. Quantification of MDC formation in Snf1-associated mutants following 2 h glucose restriction. C. Immunoblot analysis of Snf1-AID-FLAG strains and controls following auxin treatment. Haploid parent strains, an untagged OsTIR1 control, and diploid Snf1-AID-FLAG strains were treated with DMSO or auxin for the indicated times. FLAG was detected by immunoblot; Ponceau staining serves as a loading control. D. Quantification of MDC formation in Snf1-AID strains following auxin-induced degradation and 2 h carbon-source switching to galactose. Cells were pretreated with DMSO or auxin for 30 min before the media shift. E. Quantification of MDC formation in Snf1-AID strains following auxin treatment and 2 h under trace-glucose conditions. Cells were pretreated with DMSO or auxin for 30 min before the media shift. F. Quantification of MDC formation in Snf1-AID strains following auxin treatment and 2 h exposure to 2-deoxy-D-glucose (2-DG). Cells were pretreated with DMSO or auxin for 30 min before 2-DG treatment. G. Quantification of MDC formation in Snf1-AID strains following auxin treatment and 2 h concanamycin A (ConcA) exposure. Cells were pretreated with DMSO or auxin for 30 min before ConcA treatment. H. Quantification of MDC formation following auxin pretreatment and glucose restriction. Control (Snf1-AID-FLAG + EV) and test (Snf1-AID-FLAG + His3-OsTIR1 + *SCM4* OE) strains were grown overnight in YPAD, pretreated with DMSO or auxin for 30 min, and subjected to 2 h glucose restriction. I. Quantification of MDC formation in EV control, SCM4 OE, *snf1Δ*, and *snf1Δ* + *SCM4* OE strains following 2 h glucose restriction. Quantification and statistics. All MDC measurements were performed with n = 3 independent experiments and 100 cells scored per replicate. Error bars represent mean ± SEM. Statistical significance was assessed using Welch’s two-tailed t-test.

**Figure S4:**
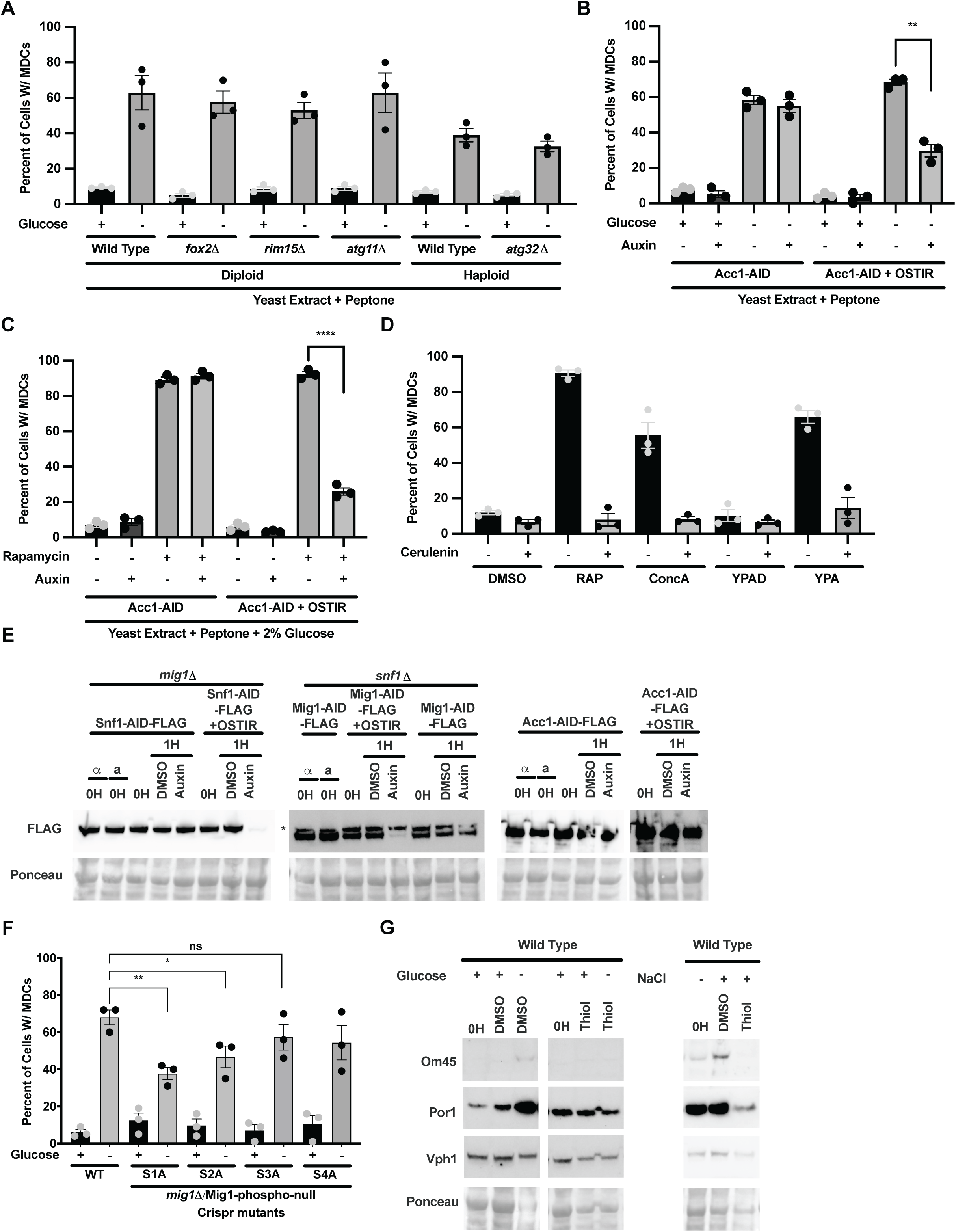
Analysis of Snf1-dependent pathways involved in MDC induction (related to Figure 4) A. Quantification of MDC formation in mutants affecting Snf1-regulated and mitochondrial remodeling pathways following 2 h glucose restriction. B. Quantification of MDC formation following auxin pretreatment and glucose restriction. Control (Acc1-AID–FLAG) and test (Acc1-AID–FLAG + His3-OsTIR1) strains were grown overnight in YPAD, pretreated with DMSO or auxin for 30 min, and subjected to 2 h glucose restriction. C. Quantification of MDC formation in Acc1-AID strains following auxin treatment and 2 h rapamycin exposure. Cells were pretreated with DMSO or auxin for 30 min before rapamycin treatment. D. Quantification of MDC formation under combined treatments with DMSO, rapamycin (RAP), concanamycin A (ConcA), YPAD (control), or YPA (glucose restriction), with or without cerulenin pretreatment. Cells were pretreated with DMSO or cerulenin for 30 min followed by 2 h treatment. E. Immunoblot analysis of AID strains used in Figure 4F and G and Figure S4B-C following auxin-induced degradation. FLAG was detected by immunoblot; Ponceau staining serves as a loading control. The asterisk indicates a nonspecific band. F. Quantification of MDC formation in CRISPR-generated Mig1 phospho-null mutants following 2 h glucose restriction. Haploid mutants were mated with *mig1Δ* strains prior to analysis. G. Immunoblot analysis of mitochondrial protein levels in wild-type cells following glucose restriction or NaCl stress in the presence of DMSO or thiolutin. Om45 and Por1 are shown; Vph1 and Ponceau staining serve as loading controls. Quantification and statistics. All MDC measurements were performed with n = 3 independent experiments and 100 cells scored per replicate. Error bars represent mean ± SEM. Statistical significance was assessed using Welch’s two-tailed t-test.

**Figure S5:**
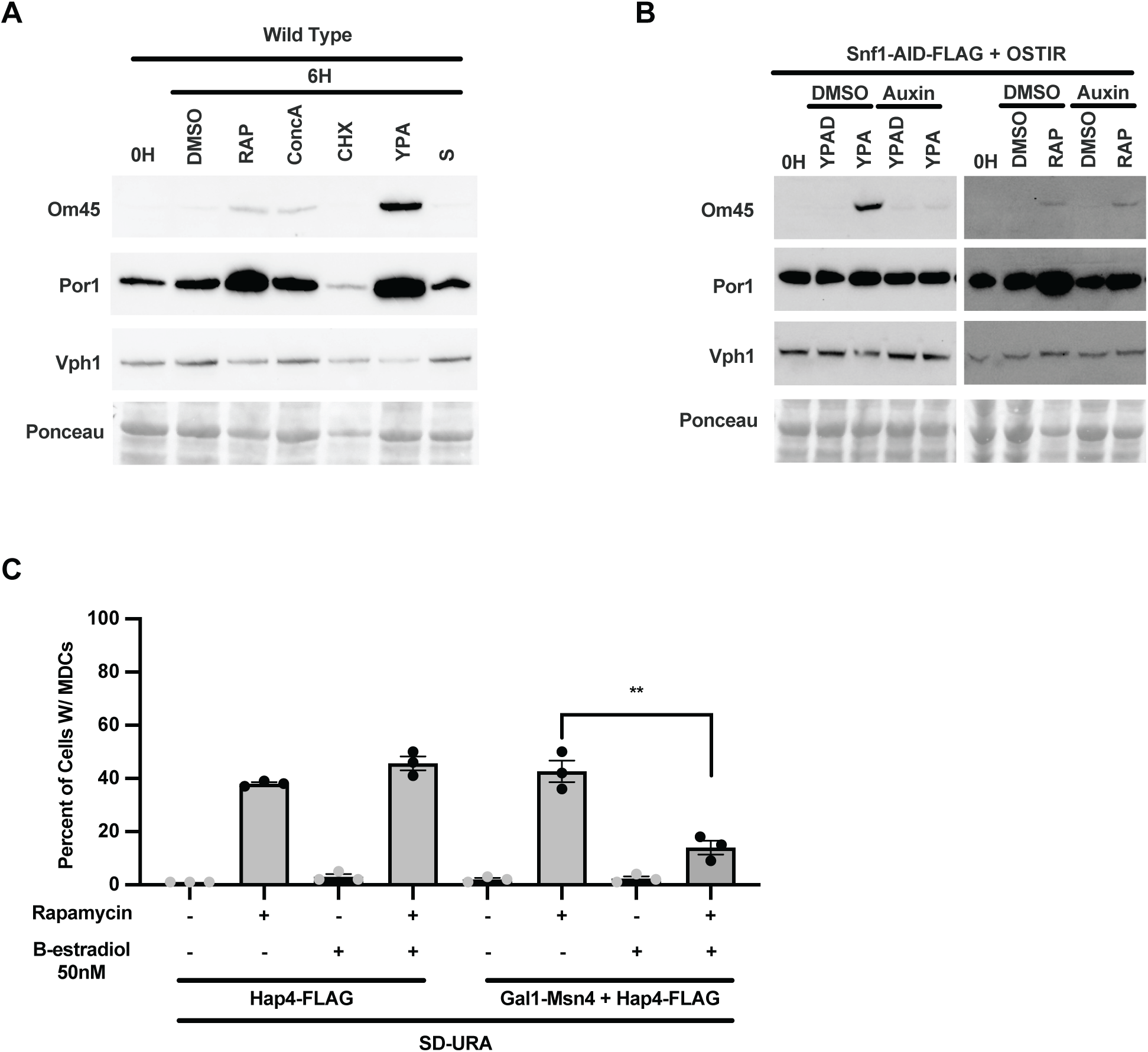
Mitochondrial protein expression and MDC regulation during distinct induction conditions (related to Figure 5) A. Immunoblot analysis of mitochondrial protein levels in wild-type cells following 6 h treatment with rapamycin, ConcA, CHX, YPA, or S. Om45 and Por1 are shown; Vph1 and Ponceau staining serve as loading controls. B. Immunoblot analysis of mitochondrial protein levels in Snf1-AID–FLAG strains following glucose restriction or rapamycin treatment with or without auxin-induced degradation. Om45 and Por1 are shown; Vph1 and Ponceau staining serve as loading controls. C. Quantification of MDC formation in Hap4 expression strains following rapamycin treatment. Cells were grown overnight in SD-URA with or without β-estradiol to induce sustained Hap4-FLAG expression and then treated with rapamycin for 2 h. Quantification and statistics. MDC measurements in (C) were performed with n = 3 independent experiments and 100 cells scored per replicate. Error bars represent mean ± SEM. Statistical significance was assessed using Welch’s two-tailed t-test.

