## Supplementary material for "Mitochondrial-derived compartments buffer outer membrane protein load during acute mitochondrial adaptation": Table S3

**Table S3. Bacterial strains, chemicals, plasmids, and software used in this study**

| Reagent or resource | Source | Identifier |
|---|---|---|
| **Antibodies** | | |
| Mouse monoclonal anti-GFP clones 7.1 and 13.1 (dilution 1:1000) | Roche | Cat # 11814460001; RRID:AB_390913 |
| Mouse monoclonal anti-PGK1 clone 22C5D8 (dilution 1:5000) | Abcam | Cat # ab113687; RRID:AB_10861977 |
| Mouse monoclonal anti-Flag M2 (1:1000) | Sigma-Aldrich | Cat # F1804 Source # SLCQ9255 |
| Mouse monoclonal anti-Por1 clone:16G9E6BC4 (dilution 1:1000) | Invitrogen Antibodies | Reference: 459500 |
| Rabbit polyclonal anti-Aco1 (dilution 1:1000) | Shaw Lab | N/A |
| Rabbit polyclonal anti-Cit1 (dilution 1:1000) | Shaw lab | N/A |
| Mouse monoclonal anti-Vph1 (dilution 1:1000) | Abcam | Cat # ab113683 RRID:AB_10D7A7B2 |
| Rabbit polyclonal anti-Om45 (dilution 1:1000) | Shaw lab | N/A |
| **Bacterial strains** | | |
| *Escherichia coli* DH5α | N/A | N/A |
| *S. cerevisiae* ORF collection (pDONR201/221) | Harvard Institute of Proteomics | N/A |
| One Shot™ *ccd*B Survival™ 2 T1$^{R}$ Competent Cells | Thermo Fisher | Cat#A10460 |
| **Chemicals, peptides, and recombinant proteins** | | |
| 3-Indoleacetic acid (Auxin) | Sigma-Aldrich | Cat # I3750 ; CAS # 87-51-4 |
| Casamino acids | US Biological | Cat # 0012501A; CAS # 65072-00-6 |
| Concanamycin A | Santa Cruz Biotechnology | Cat # sc-202111; CAS # 80890-47-7 |
| Concanavalin A | Sigma-Aldrich | Cat # L7647; CAS # 11028-71-0 |
| Cycloheximide | Sigma-Aldrich | Cat # C1988; CAS # 66-81-9 |

| | | |
|---|---|---|
| Dimethyl sulfoxide (DMSO) | Sigma-Aldrich | Cat # D2650; CAS # 67-68-5 |
| DTT | Gold Biotechnology | Cat # DTT10; CAS # 27565-41-9 / 3483-12-3 |
| Digitonin | Gold Biotechnology | Cat # D-180-2.5; CAS # 11024-24-1 |
| Lyticase | Sigma-Aldrich | Cat # L2524; CAS # 37340-57-1 |
| Rapamycin | LC Laboratories | Cat # R-5000; CAS # 53123-88-9 |
| 2-deoxy-D-glucose | Thermo Fisher Scientific | Cat # 111980050; CAS # 154-17-6 |
| Thiolutin | Sigma-Aldrich | Cat #T3450; CAS # 87-11-6 |
| Cerulenin | Merck Millipore | Cat # 219557 CAS # 17397-89-6 |
| Beta-estradiol | Sigma-Aldrich | Cat #E8875 CAS # 50-28-2 |
| **Critical commercial assays** | | |
| Bicinchoninic Acid Protein Assay | Thermo Fisher | Cat # 23227 |
| Gateway LR Clonase II Enzyme Mix | Thermo Fisher | Cat # 11791020 |
| Gibson Assembly Master Mix | New England Biolabs | Cat # E2611 |
| NEBridge Golden Gate Assembly Kit (BsmBI V2) | New England Biolabs | Cat # E1602S |
| **Plasmids** | | |
| Plasmid: pAG306GPD-HAP4 chr 1 | Schuler et al. 2021 | B3898 |
| Plasmid: pAG306GPD-ccdB chr 1 | Hughes and Gottschling, 2012 | B3681 |
| Plasmid: pAG306GPD-SCM4 chr 1 | This Study | B4027 |
| Plasmid: pHLUM | Mülleder *et al.*, 2012; Addgene | Plasmid # 40276 |
| Plasmid: pHYG-AID*-6FLAG | Morawska and Ulrich, 2013 Addgene | Plasmid # 99519 |
| Plasmid: pKT127-mCherry | Daniel Gottschling (Calico) | N/A |
| Plasmid: pKT128 | Sheff and Thorn, 2004; Addgene | Plasmid # 8729 |
| Plasmid: pRS305 | Sikorski and Hieter, 1989 | N/A |

| | | |
|---|---|---|
| Plasmid: pRS306 | Sikorski and Hieter, 1989 | N/A |
| Plasmid: pRS400 | Daniel Gottschling (Calico) | N/A |
| Plasmid: pRS40Hyg | Daniel Gottschling (Calico) | N/A |
| Plasmid: pJW1663 | Wiessman, Costa et al. 2018 | Addgene 112037 B3960 |
| Plasmid: pJW1666 | Wiessman, Costa et al. 2018 | Addgene 112040 B4077 |
| Plasmid: pJW-HAP4-FLAG-1666 | This study | |
| Plasmid: pM693 (NOP1pr:hCAS9::NatMX (Met2)) | Matt Miller's lab | Based off (DiCarlo *et al*, 2013; Ryan *et al*, 2014) |
| Plasmid: pM696 (CrispR-Guide:BsmBI:N20:PAM) | Matt Miller's lab | Based off (DiCarlo *et al*, 2013; Ryan *et al*, 2014) |
| Plasmid: pM1525 (CrispR-guide:BsmBI-PAMdonor:URA) | Matt Miller's lab | Based off (DiCarlo *et al*, 2013; Ryan *et al*, 2014) |
| Plasmid: pM1525-mig1-A-PAM-S222A+ S278A | This Study | |
| Plasmid: pM1525-mig1-B-PAM-S311A+ S381A | This Study | |
| Plasmid: pM696-Mig1-PAM222 | This Study | |
| Plasmid: pM696-Mig1-PAM278 | This Study | |
| Plasmid: pM696-Mig1-PAM311 | This Study | |
| Plasmid: pM696-Mig1-PAM381 | This Study | |
| **Software and algorithms** | | |
| FIJI | Schindelin *et al*., 2012 | Version 1 |
| Prism | GraphPad Software, Inc. | Version 9 |
| SnapGene | GSL Biotech | Version 4.2 |
| ZEN Blue Edition | Carl Zeiss Microscopy | Version 2.6 |
